# Peroxiredoxin 6 limits mitochondrial peroxidation to prevent mitochondrial ER contact site assembly and inflammatory signalling following adaptive stress

**DOI:** 10.64898/2026.07.24.740509

**Authors:** Penglin Li, Yating Zheng, Jiwang Tang, Qin Xia, Jose C. Casas-Martinez, Ángel Ortiz-Alcántara, Raquel Requejo-Aguilar, C. Alicia Padilla, Leo R. Quinlan, Antonio Miranda Vizuete, Katarzyna Goljanek-Whysall, Brian McDonagh

## Abstract

Peroxiredoxin 6 (PRDX6) is a multifunctional enzyme with peroxidase, calcium independent phospholipase A_2_ (aiPLA₂) and lysophospatidylcholine acyltransterase (LPCAT) activities. Although PRDX6 can repair peroxidised phospholipids and help prevent ferroptosis, its role in the adaptive response to physiological oxidative stress remains unclear. In this study PRDX6 function was investigated using myoblasts exposed to an acute low concentration of H_2_O_2_ and *Caenorhabditis elegans* subjected to a swimming intervention. Mild oxidative stress promoted myogenesis and mitochondrial turnover in myoblasts, while exercise enhanced activity and longevity in *C. elegans*. The adaptive responses were associated with increased mitochondrial localisation of PRDX6. In contrast, loss of PRDX6 under mild stress conditions resulted in increased mitochondrial lipid peroxidation, enhanced mitochondrial ER contact sites (MERCS), mitochondrial calcium accumulation, release of mitochondrial DNA and activation of innate immune signalling pathways. Similar phenotypes were induced by the ferroptosis activator erastin and rescued by the lipid peroxyl scavenger Ferrostatin-1, indicating lipid peroxidation was the key triggering event. Furthermore, inhibition of mitochondrial calcium uptake in *C. elegans* prevented calcium overload and attenuated inflammatory signalling. Together, the results identify PRDX6 as a conserved regulator of mitochondrial adaptation to physiological oxidative stress, functioning to limit mitochondrial lipid peroxidation and prevent excessive MERCS assembly, mitochondrial calcium dysregulation and inflammatory activation.

## Introduction

Ferroptosis is a distinct form of regulated cell death that occurs as a result of iron-catalysed accumulation of peroxidised PUFA-containing phospholipids and loss of membrane integrity [1]. Oxidation of phospholipids can create a self-propagating cascade of membrane oxidation, with accumulation of oxidised lipids due to a failure to repair membrane lipids and overwhelming cellular defence systems. Ferroptosis depends on ferrous iron (Fe²⁺) which initiates lipid peroxidation cascades by the Fenton reaction and generation of the highly reactive hydroxyl radical [2]. Mitochondria are the most active intracellular sites for iron utilisation, including haem synthesis and iron-sulphur cluster assembly, and they also store iron through mitochondrial ferritin [1]. Unlike apoptosis or necrosis, ferroptosis is characterised morphologically by reduced mitochondrial volume, increased membrane density, and cristae loss, while the nucleus remains relatively intact, indicating that mitochondria are among the earliest organelles affected during the initiation and progression of ferroptosis [1, 3]. As mitochondria cannot synthesise some of the major phospholipid classes de novo, such as phosphatidylcholine, phosphatidylinositol, and sphingolipids, they depend heavily on phospholipid precursors supplied by lipid exchange from the ER via mitochondrial ER contact sites (MERCS) [4]. Mitochondrial specific cardiolipin is particularly sensitive to peroxidation as it contains a high proportion of PUFAs and enriched at the inner mitochondrial membrane in close proximity to the components of electron transport chain. Cardiolipin is primarily synthesised from phosphatidic acid that is imported into the mitochondria from the ER via MERCS [5].

Cellular homeostasis is maintained by inter-organelle crosstalk and the exchange of material and information in response to biological perturbations. Mitochondria and ER are key regulatory hubs for maintaining cellular homeostasis and have a synergistic relationship influencing their function and adaptability to changes in the cellular environment [6]. MERCS are crucial for effective crosstalk between these two organelles, regulating iron homeostasis, metabolite exchange such as Ca^2+^ and lipids, as well as determining the sites of mitochondrial fission and fusion [7–10]. Furthermore, the initiation of ferroptosis in cells triggers increased mitochondrial fission with the recruitment of DRP1 [11]. MERCS are relatively stable structures between the surfaces of mitochondria and the ER and the width of the contact site can determine the preference for the exchange of metabolites (30 - 50 nm for Ca^2+^ and < 10 nm for lipid signalling) [7]. In pathophysiological conditions, disrupted mitochondria and ER communication result in resistance to mitochondrial degradation, accumulation of dysfunctional mitochondria, increased susceptibility to ferroptosis, release of proinflammatory mtDNA and an amplification of the pathophysiological response [6, 12]. As MERCS regulate Ca^2+^ homeostasis, redox signalling and lipid transfer, they are signalling hubs that can modulate mitochondrial dynamics, protein homeostasis and inflammation [13, 14]. The disruption of MERCS assembly/disassembly plays a major role in pathophysiological conditions particularly in age-related diseases [8]. Moreover, a recent study highlighted that MERCs are primary intracellular hotspots where phospholipid peroxidation begins during ferroptosis [15]. Lipid peroxides first accumulate at these contact sites and then propagate to mitochondria, triggering mitochondrial ROS and mitochondrial fission [15]. Although it has also been suggested that the onset of lipid peroxidation is at ER-Golgi vesicles which is transmitted to other organelles such as mitochondria and lysosomes [16].

A disruption in the intracellular redox environment can result in lipid peroxidation and ferroptosis signalling. The selenocysteine containing Glutathione peroxidase 4 (GPX4) was identified as the principal enzyme involved in the repair of oxidised phospholipids and prevention of ferroptosis [17]. However, it was previously suggested that GPX4 and Peroxiredoxin 6 (PRDX6) are the only enzymes identified to be able to reduce phospholipid hydroperoxides at a significant rate and are key for membrane repair [18, 19]. Moreover, the role of PRDX6 in prevention of redox dependent ferroptosis has been identified [19–22]. Peroxiredoxins (PRDX’s) are a family of highly conserved and abundant antioxidant enzymes that have a fundamental role in regulating cellular redox homeostasis. The PRDX family include typical 2-Cys PRDX’s 1-4, the atypical 2-Cys PRDX5 and the 1-Cys PRDX6. Together, PRDX’s make up about ∼1% of total cellular protein and family members differ in subcellular localisation and substrate specificity [23]. PRDX6 has been identified as the most conserved member of the PRDX family across 18 metazoan species analysed [24]. PRDX6 is unique as it contains a conserved peroxidatic Cys (Cys47) but lacks a resolving Cys residue, although it can use GSH with NADPH or glutathione S-transferase π to reduce its peroxidatic Cys during the catalytic cycle [18]. PRDX6 also has multifunctional enzymatic capabilities, including peroxidase activity, calcium independent phospholipase A_2_ (aiPLA₂) and lysophospatidylcholine acyltransterase (LPCAT) activities [18, 25]. The role of PRDX6 in lipid remodelling has been increasingly recognised, it can reduce phospholipid hydroperoxides (PL-OOH) to alcohols, while aiPLA₂ and LPCAT activities replace oxidised sn-2 fatty acyl groups and regenerates structurally intact, reduced phospholipids [18, 25, 26].

Global lipidomic analyses of PRDX6-deficient cells highlight its role in phospholipid homeostasis, PRDX6 loss results in reductions in sphingomyelin and acylcarnitine species, dysregulated plasmalogen homeostasis, and reduced polyunsaturated fatty acid-containing phospholipids across several glycerophospholipid classes [21]. Due to its enzymatic activities PRDX6 has an important role in ferroptosis regulation. It acts as a suppressor of phospholipid peroxide-driven cell death in parallel with GPX4 [19]. Single-point mutation studies further show that both peroxidase activity (Cys47) and aiPLA₂ activity (Asp140) are required for full ferroptosis resistance [22]. Moreover, PRDX6 can facilitate the membrane translocation of selenocysteine containing GPX4 by the formation of disulphide bonds with its peroxidatic Cys47 [19]. Loss of PRDX6 sensitises cells to ferroptosis inducers such as erastin through lipid hydroperoxide accumulation [21, 22]. While increased levels of PRDX6 have been associated with increased tumour growth by inhibition of lipid peroxidation and protection from ferroptosis induced cell death [19]. Furthermore, knockout models of PRDX6 have reported mitochondrial dysfunction [27–29]. Previously PRDX6 has been identified in mitochondrial fractions, although it lacks a canonical mitochondrial targeting sequence [28, 30]. Furthermore, an age-related decline in mitochondrial PRDX6 has been reported in skeletal muscle accompanied by increased lipid peroxidation and mitochondrial electron leakage [28].

In this study, the role of PRDX6 was investigated in the adaptation of myoblasts to low concentrations of H_2_O_2_ and the effects of an exercise intervention in the model organism *Caenorhabditis elegans*, that has previously been demonstrated to improve overall healthspan. The results demonstrate that under control conditions both an acute dose of H_2_O_2_ in myoblasts and swimming intervention in *C. elegans* promotes mitochondrial turnover. In myoblasts this was associated with mitochondrial localisation of PRDX6, improved myogenesis and increased stress resistance and longevity in *C. elegans*. However, in the absence of PRDX6, the acute dose of H_2_O_2_ in myoblasts or an exercise intervention in *C. elegans* lacking PRDX-6 resulted in detrimental effects as a result of initiation of ferroptosis signalling pathways. Mechanistically, this was due to increased mitochondrial lipid peroxidation, the formation of tight contact sites between the ER and mitochondria that resulted in mitochondrial calcium overload and release of mtDNA stimulating an inflammatory response. The results highlight a previously unrecognised role of PRDX6 in translocating to mitochondria to prevent mitochondrial lipid peroxidation, ferroptosis signalling and subsequent inflammatory signalling.

## Results

### Acute H_2_O_2_ treatment of myoblasts improves myogenesis and mitochondrial turnover that requires PRDX6

It has previously been demonstrated that acute low concentrations of H_2_O_2_ can alter myogenesis and mitochondrial dynamics [31, 32]. C2C12 myoblasts were treated for 10 min with a range of H_2_O_2_ concentrations (5-100 µM), media was replaced with fresh growth media (GM) for 24h, exchanged for differentiation media (DM) and cells were allowed to differentiate into myotubes for 7 days. Acute treatment with 25 µM H_2_O_2_ improved myogenesis as assessed by the % myotube area, myotube area and fusion index and this concentration was selected for further studies (Suppl. Fig. 1A). In this study the role of PRDX6 in the adaptive response to physiologically relevant concentration of H_2_O_2_ was determined. Following siRNA mediated knockdown of *Prdx6* and treatment with H_2_O_2_ (Suppl. Fig. 1B), the levels of PRDX1, PRDX2, PRDX3 and PRDX5 did not change indicating no compensatory changes in levels of these proteins (Suppl. Fig. 1C-F). The improved myogenesis following acute H_2_O_2_ treatment was clear but not detected following knockdown of *Prdx6*, which had decreased myotube area and diameter (Fig. 1A). Although it was clear that knockdown of *Prdx6* alone had significant effects on myogenesis, with decreased myotube area and diameter compared to controls (Fig. 1A). The adaptive response of myoblasts to an acute treatment of H_2_O_2_ resulted in increased mitochondrial content and dynamics [31]. Following the 10 min H_2_O_2_ treatment cells were allowed to recover in growth media and there were increased levels of the mitochondrial marker TOMM20, increased TFAM a key regulator of mitochondrial DNA synthesis and mitochondrial associated proteins Protein DJ-1 and VDAC1, but these changes were not detected following knockdown of *Prdx6* (Suppl. Fig. 1G-J). Interestingly no change in the levels of PGC1α were apparent (Suppl. Fig. 1K). After the recovery period following the acute bolus addition of 25 µM H_2_O_2_ increased mitochondrial content as assessed by MitoTracker green with no increase in mitochondrial ROS generation assessed by MitoSOX staining (Fig. 1B,E,F), TMRM staining demonstrated increased mitochondrial membrane potential (Fig. 1C,G). Mitophagy was determined using the mitochondrial targeted Cox8-EGFP-mCherry reporter construct, a decreased GFP:RFP ratio indicated increased mitophagy following H_2_O_2_ treatment (Fig. 1D,H). However, these changes were not detected following knockdown of *Prdx6*, although there was increased MitoSOX staining following H_2_O_2_ treatment (Fig. 1B-D). The increased mitophagy in control cells treated with H_2_O_2_ was accompanied by an increased ratio of LC3 II/I, increased levels of ULK1 and decreased levels of p62, suggesting increased autophagic flux (Suppl. Fig1L-N). Similarly, qPCR analysis of mitophagy related genes, *Pink1*, *Parkin*, *Bnip3* and *Becn1*, revealed increased levels following H_2_O_2_ treatment that was not apparent following knockdown of *Prdx6* (Suppl. Fig. 1O). Furthermore, 25 µM H_2_O_2_ treatment resulted in increased filamentous mitochondria with increased levels of OPA1 and MFN2, while loss of PRDX6 resulted in increased mitochondrial fragmentation and elevated levels of DRP1 (Suppl. Fig. 2A-D) Interestingly following 25 µM H_2_O_2_ treatment, cytoplasmic and mitochondrial fractionation revealed a large increase in mitochondrially localised PRDX6 (Fig. 1I). This was further supported by immunofluorescence analysis (Fig. 1J). Together these results demonstrate that an acute 25 µM H_2_O_2_ treatment resulted in PRDX6 mitochondrial translocation, increased mitochondrial content and promoted autophagy in myoblasts that ultimately resulted in improved myogenesis, which was not apparent in myoblasts with siRNA knockdown of *Prdx6*.

**Fig. 1.**
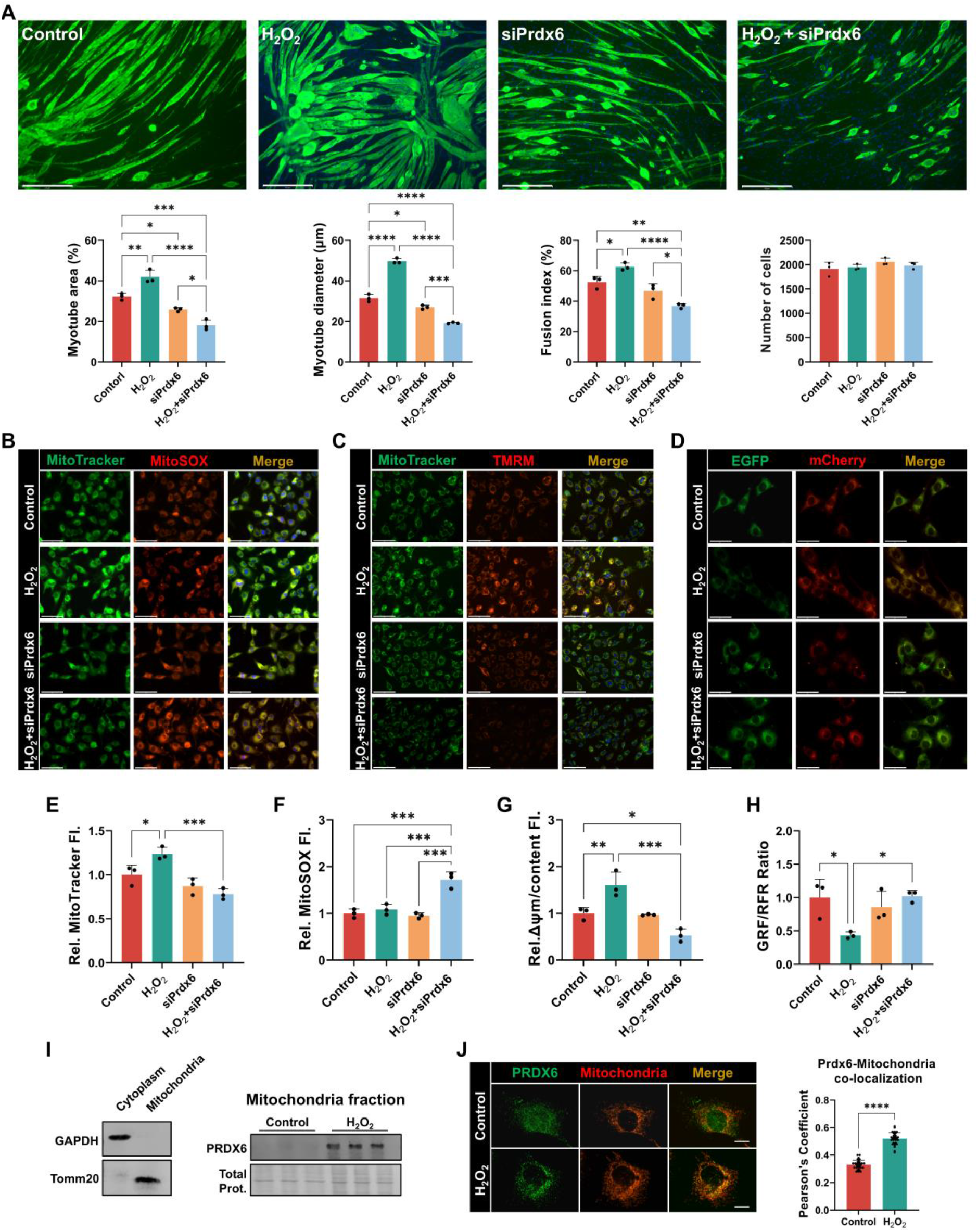
Acute H_2_O_2_ promotes myogenesis and mitochondrial turnover that requires PRDX6 in C2C12 myoblasts. **(A)** Representative immunofluorescence images and quantification of myotubes after 7 days of differentiation. Myoblasts were transfected with control or *Prdx6* siRNA, treated for 10 min with 25 µM H_2_O_2_ or vehicle, recovered in growth medium for 24 h and then differentiated. Myotubes were labelled with the MF20 antibody (myosin heavy chain) and nuclei counterstained with DAPI. Graphs represent myotube area, myotube diameter, fusion index and total cell number. Scale bars, 275 µm. **(B-D)** Representative images of myoblasts after the 10 min treatment and recovery period. Mitochondrial content (MitoTracker Green) and mitoROS (MitoSOX), scale bars, 75 µm (B). Mitochondrial content (MitoTracker Green) and membrane potential (TMRM), scale bars, 75 µm (C). Mitophagy assessed with the mitochondrially targeted Cox8-EGFP-mCherry reporter, including EGFP and mCherry signals, scale bars, 50 µm (D). **(E-H)** Quantification of relative MitoTracker fluorescence (E), relative MitoSOX fluorescence (F); mitochondrial membrane potential normalised to mitochondrial content (TMRM relative to MitoTracker) (G), and ratio of EGFP to mCherry fluorescence of the Cox8-EGFP-mCherry reporter, where a lower ratio indicates increased mitophagy (H). **(I)** Immunoblots of cytoplasmic and mitochondrial fractions. GAPDH and TOMM20 confirm fraction purity. PRDX6 in the mitochondrial fraction of control and H_2_O_2_ treated cells, with total protein as the loading control. **(J)** Representative images and Pearson’s correlation coefficient for co-localisation of PRDX6 with mitochondria in control and H_2_O_2_ treated cells. Scale bars, 50 µm. Data presented as mean ± SEM. For A-H, n =3, statistical significance was determined by two-way ANOVA. For J, n=30, statistical significance was determined by student’s t-test. \**p* < 0.05, ** *p* < 0.01, *** *p* < 0.001, **** *p* < 0.0001.

### Loss of PRDX6 phospholipase activity results in lipid peroxidation and ferroptosis following H_2_O_2_ treatment

PRDX6 is a multifunctional enzyme with both peroxidase and phospholipase A2 activities and its role in the prevention of ferroptosis has recently been identified [25]. In order to dissect the different functional roles of PRDX6 following H_2_O_2_ treatment in the myoblast model, MJ33 was included to specifically inhibit the phospholipase A2 activity of PRDX6 [33]. Furthermore, as PRDX6 has been identified as a key regulator of ferroptosis, treatment with the ferroptosis inducer, erastin, was included together with a scavenger of lipid peroxyl radicals that can inhibit ferroptosis, Ferostatin-1 (Fer-1). Consistently, acute treatment with H_2_O_2_ resulted in increased myotube area and diameter, while knockdown of *Prdx6* resulted in decreased myotube area and diameter (Fig. 2A). However, treatment of cells with MJ33 also resulted in decreased myotube area and diameter, indicating the phospholipase A2 activity of PRDX6 was essential for this adaptive response. Moreover, these results were similar to the decreased myogenesis following treatment of myoblasts with erastin. Furthermore, Fer-1 rescued this phenotype when combined with *Prdx6* knockdown, MJ33 or erastin (Fig. 2A). These results highlight that the phospholipase A2 activity of PRDX6 was essential during myogenesis following a relatively low and acute treatment with 25 µM H_2_O_2_, and inhibition of phospholipases activity results in cells undergoing ferroptosis. As lipid peroxidation is the key driver of ferroptosis signalling, Bodipy C11 staining was performed to assess overall lipid peroxidation (Fig. 2B) together with co-staining of mitochondria MitoTracker Deep Red to determine if mitochondrial lipid peroxidation was affected (Fig. 2C). Consistently a combination of H_2_O_2_ with either loss of PRDX6 or inhibition of its phospholipase activity with MJ33 resulted in increased lipid peroxidation that was predominantly localised to mitochondria, similar to what was observed with erastin treatment. The increased lipid peroxidation could be rescued under those conditions with Fer-1 treatment (Fig. 2B,C). To confirm the changes detected by Bodipy C11 staining, total and mitochondrial levels of malondialdehyde (MDA), glutathione (GSH) and GPX4 levels were determined. The knockdown of *Prdx6* or inhibition of its phospholipase A2 activity with MJ33 in combination with H_2_O_2_ resulted in increased overall MDA and decreased GSH levels (Fig. 2D,E) and in mitochondrial fractions (Fig. 2F,G), comparable to treatment with erastin. Fer-1 could rescue this phenotype suggesting it was ferroptosis dependent (Fig. 2D-G). Similarly, under ferroptosis conditions there was decreased levels of overall and mitochondrially localised GPX4, which could be rescued with Fer-1 (Fig. 2H,I). Together, these results highlight that under mild oxidative conditions that can promote myogenesis, inhibition of PRDX6 phospholipase A2 activity results in increased mitochondrial lipid peroxidation and ferroptosis.

**Fig. 2.**
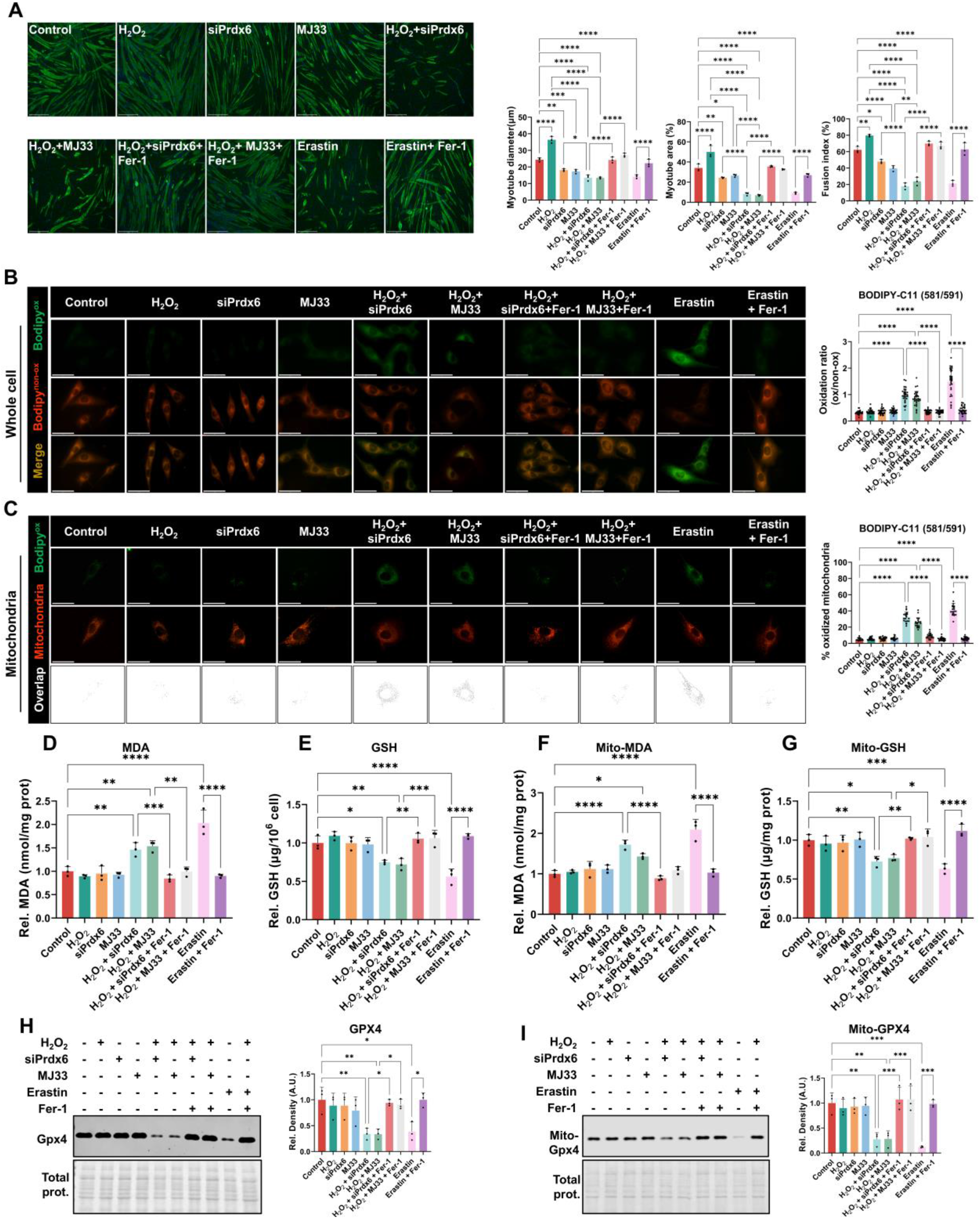
Loss of PRDX6 phospholipase A2 activity result in mitochondrial lipid peroxidation and ferroptosis phenotype following H_2_O_2_ treatment. Myoblasts were transfected with control or *Prdx6* siRNA, or treated with the PRDX6 phospholipase A2 inhibitor MJ33, the ferroptosis inducer erastin, or the lipid radical scavenger ferrostatin-1 (Fer-1), alone or combined with a 10 min treatment with 25 µM H_2_O_2_. **(A)** Representative immunofluorescence images and quantification of myotubes after 7 days of differentiation, labelled with the MF20 antibody and counterstained with DAPI. Graphs represent myotube diameter, myotube area and fusion index. Scale bars, 275 µm. **(B)** Whole cell lipid peroxidation assessed by BODIPY-C11 (581/591), showing the oxidised (green) and non-oxidised (red) signals and the merge. Quantification shows the oxidised to non-oxidised ratio. Scale bars, 50 µm. (**C)** Mitochondrial lipid peroxidation assessed by BODIPY-C11 (581/591, oxidised, green) co-stained with MitoTracker Deep Red. The overlap demonstrates the oxidised signal within mitochondria. Quantification shows the percentage of oxidised mitochondria. Scale bars, 50 µm. (**D-G) (D)** Quantification of total malondialdehyde (MDA), **(E)** total glutathione (GSH), **(F)** mitochondrial MDA and **(G)** mitochondrial GSH. **(H, I)** Immunoblots and quantification of total GPX4 and mitochondrial GPX4, with the treatment matrix indicated above each blot. Total protein was used as the loading control. Data presented as mean ± SEM. For A, D-I, n = 3; for B, n = 30; for C, n = 20. Statistical significance was determined by one-way ANOVA with Tukey’s multiple comparisons test. * *p* < 0.05, ** *p* < 0.01, *** *p* < 0.001, **** *p* < 0.0001.

### Loss of PRDX6 phospholipase A2 activity results in increased MERCS assembly and large influx of mitochondrial calcium

It was recently demonstrated that MERCS are key sites for the lipid peroxidation involved in driving ferroptosis [15]. As our results have demonstrated that PRDX6 localises to mitochondria under mild oxidative conditions and loss of its phospholipase A activity results in mitochondrial lipid peroxidation and ferroptosis, a split-FAST reporter system was used to determine the effects on MERCS assembly [34] (Fig. 3A). Similar to the lipid peroxidation results, a combination of H_2_O_2_ with either loss of PRDX6 or MJ33 treatment resulted in increased MERCS formation that could be rescued with Fer-1, consistent with erastin treatment (Fig. 3B,C). As MERCS are crucial for not only the exchange of lipids between the ER and mitochondria but also for channelling Ca^2+^ between these organelles, Rhod-2 staining was used to determine the effects on mitochondrial Ca^2+^ levels and Fluo-4 was used to assess changes in cytoplasmic Ca^2+^ levels. No changes in cytoplasmic Ca^2+^ levels were detected apart from when the positive control Thapsigargin was included (Suppl. Fig3). Co-staining of mitochondria with MitoTracker green and Rhod-2 revealed that conditions that promoted increased MERCS formation such as knockdown of PRDX6 or MJ33 in combination with H_2_O_2_, resulted in a large increase in mitochondrial Ca^2+^ levels similar to erastin treatment and could be rescued with Fer-1 (Fig. 3D,E). As a result of the increased MERCS formation and mitochondrial Ca^2+^ levels, qPCR analysis of genes for Ca^2+^ channels in mitochondria (mitochondrial calcium uniporter, Mcu) and the ER/SR (Inositol 1,4,5-trisphosphate-gated calcium channel (Itpr1), sarcoplasmic/endoplasmic reticulum ATPase 2 (Atp2a2), ryanodine receptor 1 (Ryr1)) along with calcium/calmodulin dependent kinase 1d (Camk1d) were analysed. Under the same conditions that increased mitochondrial Ca^2+^ levels, lipid peroxidation and MERCS formation, there was increased levels of *Mcu*, *Itpr1*, *Atp2a2* and *Camk1d*, while *Ryr1* levels only increased following erastin treatment (Fig. 3F-J). Together the results highlight that increased MERCS as a result of loss of PRDX6 phospholipase activity in mild oxidative stress conditions or erastin treatment, resulted in increased mitochondrial Ca^2+^ and increased levels of genes involved in Ca^2+^ transport.

**Fig. 3.**
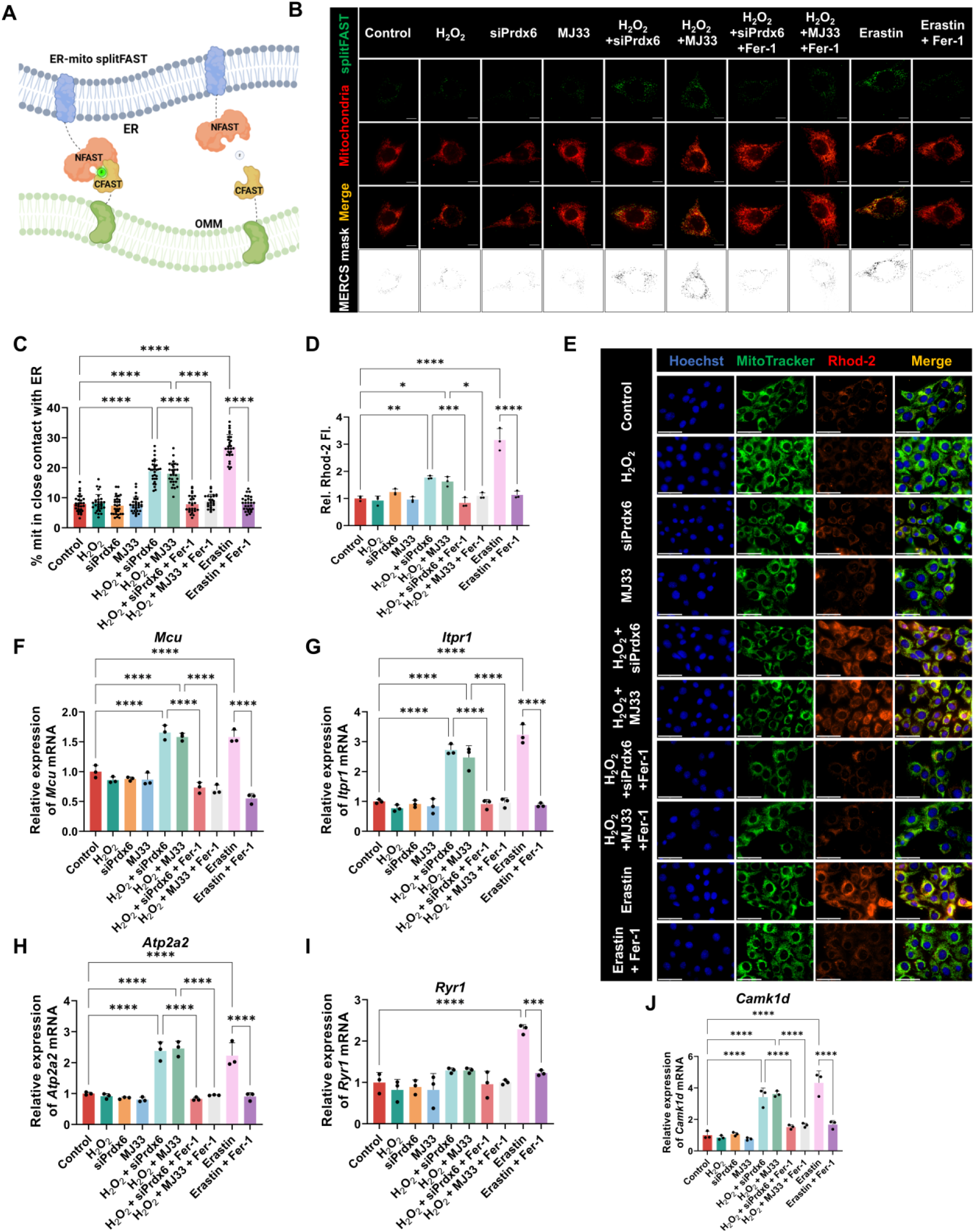
Loss of PRDX6 phospholipase A2 activity increases MERCS assembly and mitochondrial calcium influx. **(A)** Schematic of the ER-mitochondria splitFAST reporter, in which the NFAST and CFAST fragments are targeted to the ER membrane and the outer mitochondrial membrane and reconstitute a fluorescent signal at contact sites. **(B)** Representative images of the splitFAST signal, mitochondria and the merged images, with the derived MERCS mask shown below. Scale bars, 10 µm. (**C)** Quantification of the percentage of mitochondria in close contact with the ER. (**D)** Quantification of relative mitochondrial Rhod-2 fluorescence**. (E)** Representative images of myoblasts stained with Hoechst, MitoTracker Green and Rhod-2 with the merged image. Scale bars, 50 µm. **(F-J)** Relative mRNA expression levels of the calcium channel genes *Mcu*, *Itpr1*, *Atp2a2*, *Ryr1* and *Camk1d*. Data presented as mean ± SEM. For C, n = 26-30; for D and F-J, n = 3. Statistical significance was determined by one-way ANOVA with Tukey’s multiple comparisons test. * *p* < 0.05, ** *p* < 0.01, *** *p* < 0.001, **** *p* < 0.0001.

### Increased mitochondrial Ca^2+^ results in mtDNA release and cGAS-STING mediated inflammatory signalling

Sustained or excessive mitochondrial Ca^2+^ can result in opening of the mitochondrial transition pore (mPTP) and initiation of pro-apoptotic signalling [35, 36]. One of the key signalling responses following mPTP opening and loss of mitochondrial membrane potential is the release of mtDNA into the cytoplasm. Initially an immunostaining approach with anti-DNA and MitoTracker Deep Red revealed that there was a significant increase in extramitochondrial DNA defined as DNA puncta that did not overlap with the mitochondrial stain under conditions of increased mitochondrial Ca^2+^ and MERCS formation, H_2_O_2_ treatment with knockdown of *Prdx6* or inclusion of MJ33, similar to what was observed with erastin treatment (Fig. 4A). To confirm the release of mtDNA, a qPCR approach for mitochondrial genes in cytoplasmic fractions was performed. The levels of mtDNA encoded genes *Cytb*, *nd1*, *D-loop* and *Rnr2* all increased in cytoplasmic fractions as determined by qPCR in conditions with increased mitochondrial Ca^2+^ (Fig. 4B-E). Due to its ancestral bacterial origin, mitochondrial DNA is known as a damage associated molecule pattern (DAMP) that can stimulate an inflammatory response in mammalian cells. Due to the increased mtDNA release following inhibition of PRDX6 phospholipase activity with MJ33 treatment combined with H_2_O_2_, we analysed a range of inflammatory response signals. Western blot analysis of levels of cGAS, STING and Antiviral innate immune response effector (IFIT1) highlighted that under conditions of mtDNA release there was a robust increase in the cGAS/STING inflammatory response (Fig. 4F). Furthermore, qPCR analysis of genes involved in the inflammatory response were also quantified including *Ifit1*, interferon induced protein 44 (*Ifi44*), C-X-C motif 10 (*Cxcl10*) and interleukin 18 (*Il-18*). Consistently loss of PRDX6 or its phospholipase activity together with H_2_O_2_ resulted in a robust increase in inflammation associated genes similar to erastin (Fig. 4G-J). Together these results highlight under conditions that promote ferroptosis including loss of PRDX6 phospholipase A2 activity that promotes increased MERCS and excessive mitochondrial Ca^2+^, results in increased mtDNA release and markers of inflammation.

**Fig. 4.**
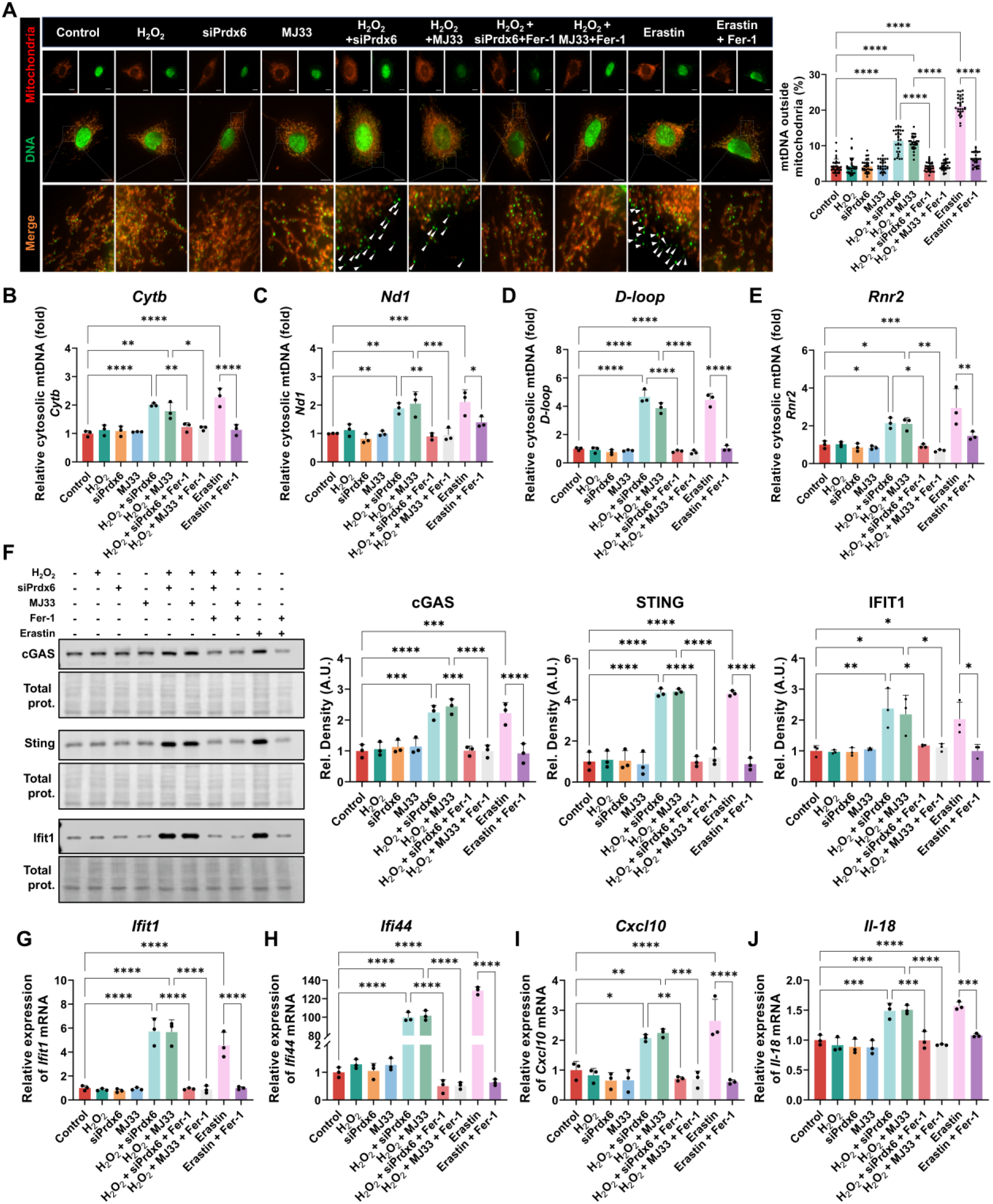
Increased mitochondrial calcium is associated with mtDNA release and cGAS-STING inflammatory signalling. **(A)** Representative immunofluorescence images of mitochondria (MitoTracker Deep Red) and DNA (anti-DNA) with the merge. Boxed regions are enlarged below and arrowheads indicate extramitochondrial DNA puncta. Quantification represents the percentage of mtDNA outside mitochondria. Scale bars, 10 µm. **(B-E)** qPCR quantification of the mitochondrial genes *Cytb*, *Nd1*, *D-loop* and *Rnr2* in cytoplasmic fractions. **(F)** Immunoblots and quantification of cGAS, STING and IFIT1, with the treatment matrix indicated above the blots. Total protein was used as the loading control. **(G-J)** Relative mRNA expression levels of the inflammatory genes *Ifit1*, *Ifi44*, *Cxcl10* and *Il18*. Data presented as mean ± SEM. For A, n = 30; for B-J, n = 3. Statistical significance was determined by one-way ANOVA with Tukey’s multiple comparisons test. * *p* < 0.05, ** *p* < 0.01, *** *p* < 0.001, **** *p* < 0.0001.

### *C. elegans prdx-6* mutant strains have impaired longevity, stress resistance and activity following an exercise intervention

In order to determine the role of PRDX6 in the context of a whole organism, we used *C. elegans* mutant strains of *prdx-6* (*tm4225* and *tm4284*) following an exercise intervention previously shown to increase stress resistance and longevity in wild type worms [31, 37]. The 5-day exercise intervention has previously been demonstrated to transiently increase endogenous ROS generation, promote mitochondrial turnover and MERCS assembly in wild type worms [31, 37]. Consistently in the N2 wild type strain, a 5-day swimming intervention resulted in increased longevity, stress resistance and indicators of physiological fitness (Fig. 5A-D). However, mutant strains lacking PRDX-6 had decreased longevity, decreased stress resistance and an altered physiological activity response following the 5-day swimming intervention (Fig. 5A-D). These results highlight that following an exercise intervention that has been demonstrated to generate an increase in intracellular ROS, there was a disrupted adaptive response in *prdx-6* mutant strains (*tm4225* and *tm4284*).

**Fig. 5.**
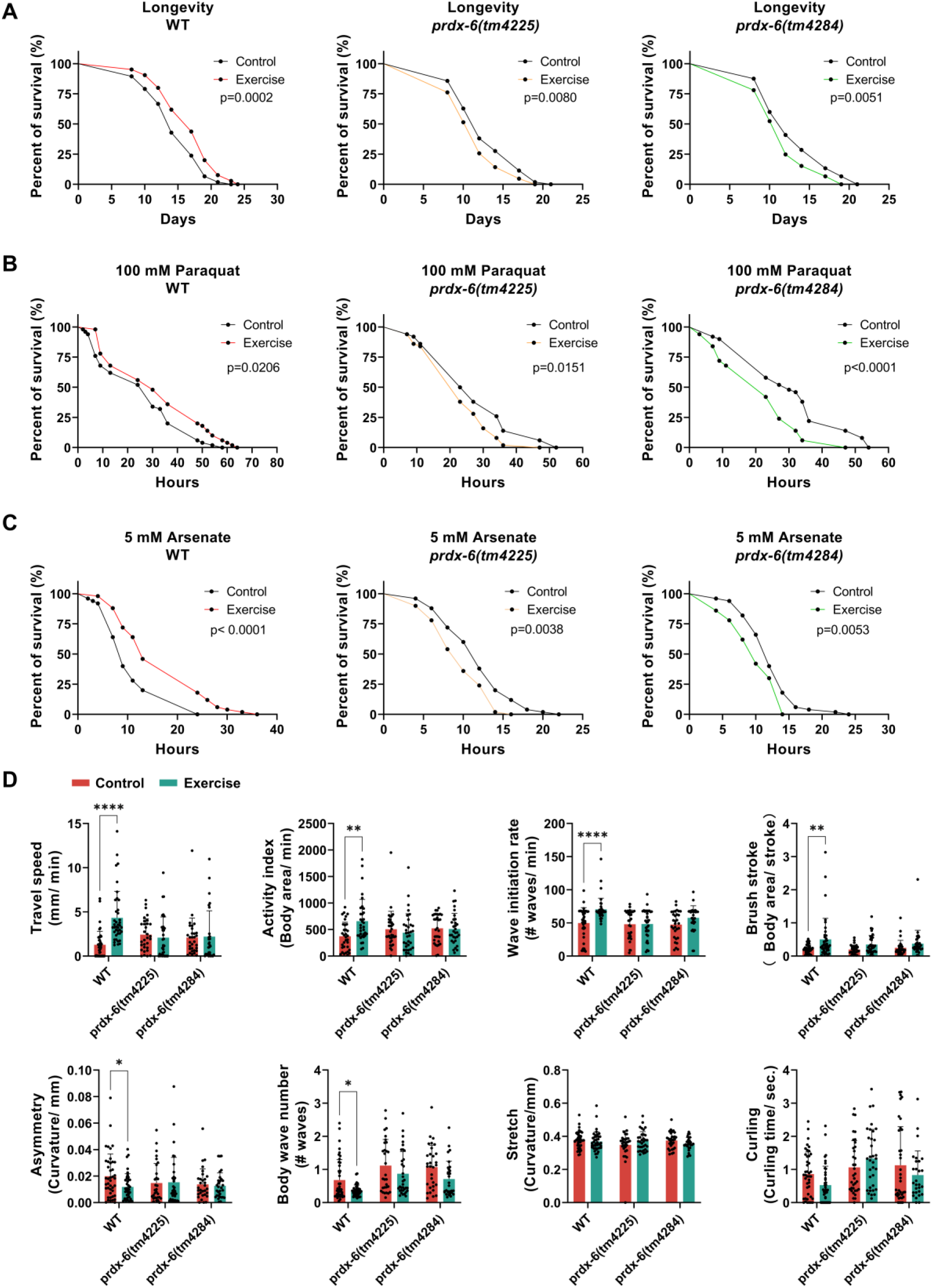
*prdx-6* mutant *C. elegans* have decreased lifespan, stress resistance and altered activity response following exercise. Wild type (N2) and *prdx-6* mutant strains *tm4225* and *tm4284* were subjected to a 5-day swimming exercise intervention and the measurements were conducted on the following day. **(A)** Kaplan-Meier survival curves for lifespan of control and exercised animals of each strain. (**B, C)** Survival curves for oxidative stress resistance following exposure to 100 mM paraquat and 5 mM arsenite. **(D)** Physiological activity (Travel speed, Activity index, Wave initiation rate, Brush stroke) and frailty (Asymmetry, Body wave number, Stretch, Curling time) monitored by CeleST in control and exercised animals of each strain. Data presented as mean ± SEM. For A-C, n = 105 and *p* values were determined by the log-rank (Mantel-Cox) test. For D, n = 30-40 and significance was determined by Student’s t-test. * *p* < 0.05, ** *p* < 0.01, **** *p* < 0.0001.

### *C. elegans prdx-6* mutants have disrupted mitochondrial dynamics following an exercise intervention

Increased mitochondrial content and dynamics have been well described in a variety of model organisms in response to exercise [38, 39]. Following the 5-day swimming intervention starting on adult Day 1, assays were performed following a recovery period on adult day 6, to investigate the changes in mitochondrial dynamics underlying the disrupted adaptation to exercise in *prdx-6* mutant strains. TMRM staining was used to assess mitochondrial membrane potential, N2 wild type worms had increased staining, while both *prdx-6* mutant strains had no change in mitochondrial membrane potential following the exercise intervention (Fig. 6A,B). A swimming intervention has previously been demonstrated to increase intracellular ROS generation immediately following exercise that returns to baseline levels after a recovery period in wild type worms [37]. On day 6 adult worms following the 5-day exercise, there was no change in mitochondrial ROS as assessed by MitoSOX staining in wild type worms, but *prdx-6* mutants had elevated MitoSOX staining following the recovery period indicating a disrupted intracellular redox environment (Fig. 6C,D). Overall mitochondrial content in the body wall muscle following the exercise was assessed using the *myo-3p*::GFP(mit) reporter on Day 6, that was crossed with *prdx-6* mutant strains (*tm4225* and *tm4284*). There was increased mitochondria in muscle of wild type worms that was not detected in the *prdx-6* mutants (Fig. 6E,F). Mitochondrial morphology can determine substrate use as mitochondrial fragmentation can increase their capacity for fatty acid oxidation [29, 40], while mitochondrial morphology changes rapidly following exercise and recovery [37]. The morphology of body wall muscle following the exercise intervention was assessed. Exercise promoted an increase in filamentous mitochondria in body wall muscle following a recovery period in the wild type strain, but *prdx-6* mutants had a distinct fragmented mitochondrial morphology in non-exercised controls and no increase in filamentous mitochondria following exercise (Fig. 6G,H). One of the regulatory determinants of mitochondrial turnover is mitophagy, the effect of exercise on mitophagy was assessed in body wall muscle using the *unc-119(ed3); Ex[myo-3p::tomm-20::Rosella; unc-119(+)]* strain that was also crossed with *prdx-6* mutant strains. In the wild type strain, the swimming intervention resulted in increased mitophagy that was not detected in either of the *prdx-6* mutants (Fig. 6I). These results highlight that exercise promoted an increase in body wall muscle mitochondrial content, increased filamentous mitochondria and mitophagy in wild type worms that requires the function of PRDX-6.

**Fig. 6.**
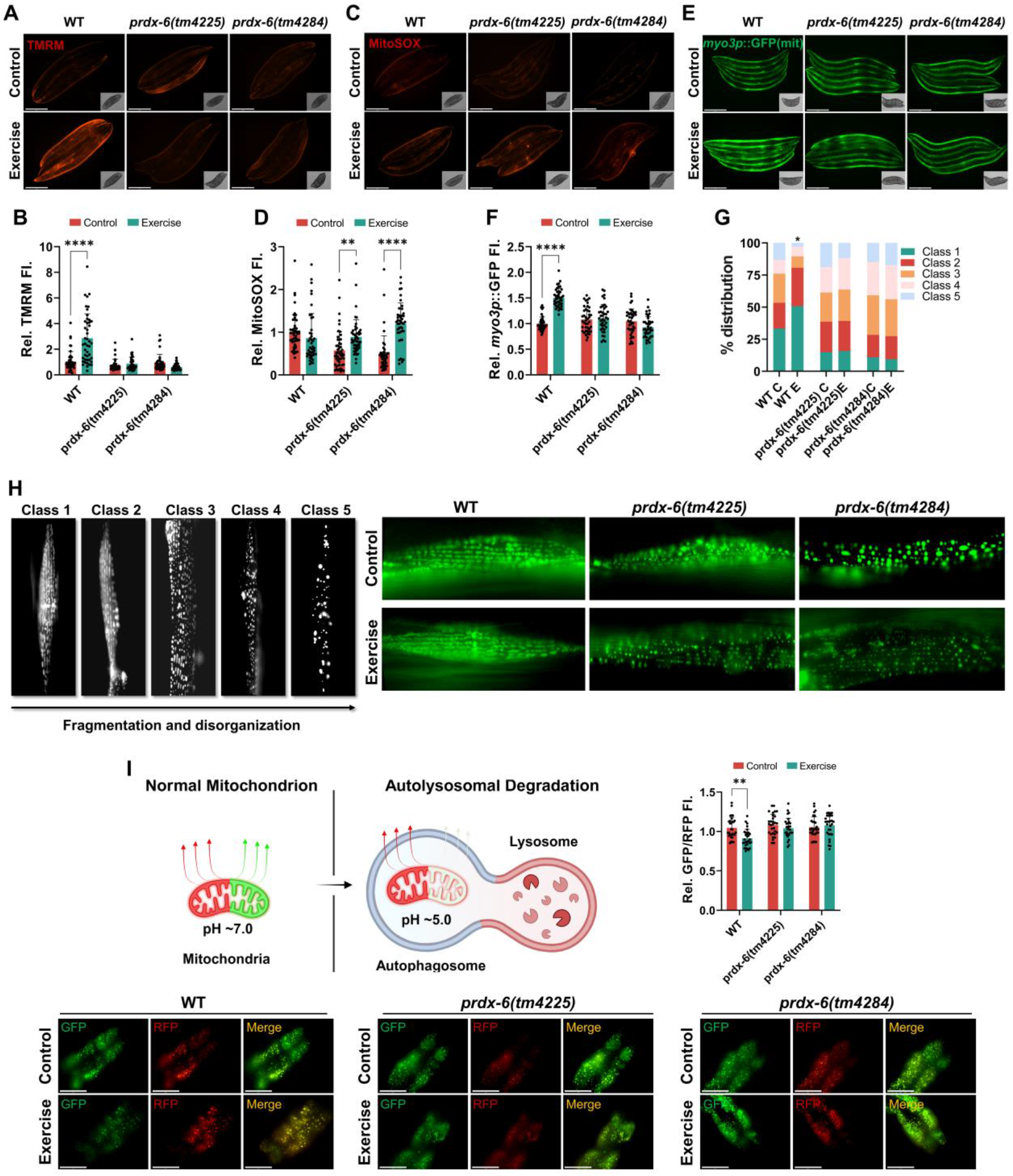
*prdx-6* mutant *C. elegans* strains have disrupted mitochondrial dynamics following exercise. Wild-type (N2) and *prdx-6* mutant strains (*tm4225* and *tm4284*) were assessed on adult day 6 following a 5-day swimming intervention or control treatment. **(A, B)** Mitochondrial membrane potential assessed by TMRM staining. **A**, representative images with brightfield insets. **B**, quantification of relative TMRM fluorescence. Scale bars, 275 µm. **(C, D)** Mitochondrial ROS assessed by MitoSOX staining, **(C)** representative images and (**D)**, quantification of relative MitoSOX fluorescence. Scale bars, 275 µm. **(E, F)** Body wall muscle mitochondrial content assessed with the *myo-3p*::GFP(mit) reporter. **(E)** representative images and **(F)** quantification of relative *myo-3p*::GFP(mit) fluorescence. Scale bars, 275 µm. **(G, H)** Body wall muscle mitochondrial morphology. **(G)** Stacked bar chart representing the percentage distribution across five morphology classes for control (C) and exercised (E) animals. **(H)** Reference images of morphology classes 1 to 5 with increasing fragmentation and disorganisation (left) and representative *myo-3p*::GFP(mit) images of body wall muscle (right). **(I)** Mitophagy assessed with the *myo-3p*::*tomm-20*::Rosella reporter. The schematic demonstrates the dual emission Rosella biosensor, in which the pH sensitive GFP signal is quenched in acidic autolysosomes, a lower GFP to RFP ratio indicates increased mitophagy. The graph shows the relative GFP to RFP ratio and representative GFP, RFP and merged images below. Scale bars, 50 µm. For B, D, F and I, data presented as mean ± SEM. Significance was determined by Student’s t-test, n = 30-40. For G, data represent the percentage distribution of 130-150 images per group, and significance was determined by chi-square test. * *p* < 0.05, ** *p* < 0.01, **** *p* < 0.0001.

### Exercise results in increased mitochondrial lipid peroxidation, increased MERCS and mitochondrial Ca^2+^ levels in *prdx-6* mutants

In the cell myoblast model knockdown of PRDX6 or inhibition of its phospholipase A2 activity with MJ33 followed by an acute 10 min treatment with 25 µM H_2_O_2_ resulted in increased mitochondrial lipid peroxidation, increased MERCS assembly and mitochondrial Ca^2+^ overload with the release of mtDNA and inflammatory signalling (Figs.2-4). Similar effects were observed when cells were treated with the ferroptosis inducer, erastin and could be rescued with the lipid peroxyl scavenger Fer-1. We investigated whether similar effects were observed in a *C. elegans* whole animal model using exercise as a physiological model or treatment with diethyl maleate (DEM) a glutathione scavenger, used to promote ferroptosis in *C. elegans* models [41, 42]. To determine lipid peroxidation Bodipy C11 staining was performed in wild type worms following exercise or DEM treatments and following exercise with the inclusion of Fer-1 in *prdx-6* mutant strains. The results highlight that DEM increased lipid peroxidation but exercise did not in the N2 wild type strain (Fig. 7A-G). In the *prdx-6* mutant strains, exercise alone was able to increase lipid peroxidation which could be prevented by the inclusion of Fer-1 (Fig. 7A-G). Furthermore, overall levels of MDA and GSH were assessed along with MDA and GSH levels in mitochondrial fractions. In wild type worms, treatment with DEM increased MDA and decreased GSH levels, which was consistent in the *prdx-6* mutant strains following exercise and could be rescued with inclusion of Fer-1 (Fig. 7C-G). To confirm the ferroptosis phenotype, qPCR was performed for a number of genes involved in the regulation of ferroptosis including *aat-9*, *ftn-1*, *gpx-1* and *acs-17* [43, 44]. The levels of genes involved in the protection against ferroptosis (*aat-9*, *ftn-1* and *gpx-1*) all decreased following DEM treatment in the wild type strain or exercise in the *prdx-6* mutants, while the levels of the pro-ferroptosis gene *acs-17* increased in those conditions (Fig. 7H). The results highlight that similar to the cell model, loss of PRDX-6 in *C. elegans* together with a physiological stress promotes ferroptosis. In order to determine whether this increased ferroptosis signature was accompanied by changes in mitochondrial associated membranes (MAMs), *vkEx2674[pnhx-2CemOrange2::PISY-1; pmyo-2GFP]; zcIs17[pges-1mitGFP]* reporter strain was used in combination with RNAi knockdown of *prdx-6* [45, 46]. DEM promoted increased MAMs in the wild type strain, while exercise increased MAMs with RNAi knockdown of *prdx-6* that could be rescued by Fer-1 (Fig. 7I and Suppl. Fig. 4A). To determine if the increased MAMs affected cytoplasmic and mitochondrial Ca^2+^ levels, we used GCaMP reporter strains *goeIs3[pmyo-3SL1::GCamP3.35::SL2; unc-119(+)];aceIs1[pmyo-3mitoLAR-GECO; pmyo-2::RFP]* and *goeIs3 [myo-3p::SL1::GCamP3.35::SL2::unc54 3’UTR + unc-119(+)]* respectively [46]. In these models DEM increased mitochondrial Ca^2+^ levels in the wild type strain and with RNAi knockdown of *prdx-6* combined with the exercise intervention (Fig. 7J and Suppl. Fig. 4A,C). No changes in cytoplasmic Ca^2+^ levels were detected under these conditions (Suppl. Fig. 4B,D) The increased MAMs and mitochondrial Ca^2+^ levels were also associated with increased expression of Ca^2+^ channels *mcu-1* (ortholog of mitochondrial calcium uniporter), *itr-1* (ortholog of Inositol 1,4,5-trisphosphate receptor), *vdac-1* (ortholog of voltage dependent anion selective channel) and *unc-68* (ortholog of ryanodine receptor) (Fig. 7K). Together these results demonstrate that similar to cell models, a physiological stress in *C. elegans* strains lacking PRDX-6 results in increased mitochondrial lipid peroxidation, increased MAMs, higher mitochondrial Ca^2+^ levels and a ferroptosis phenotype that could be rescued with Fer-1.

**Fig. 7.**
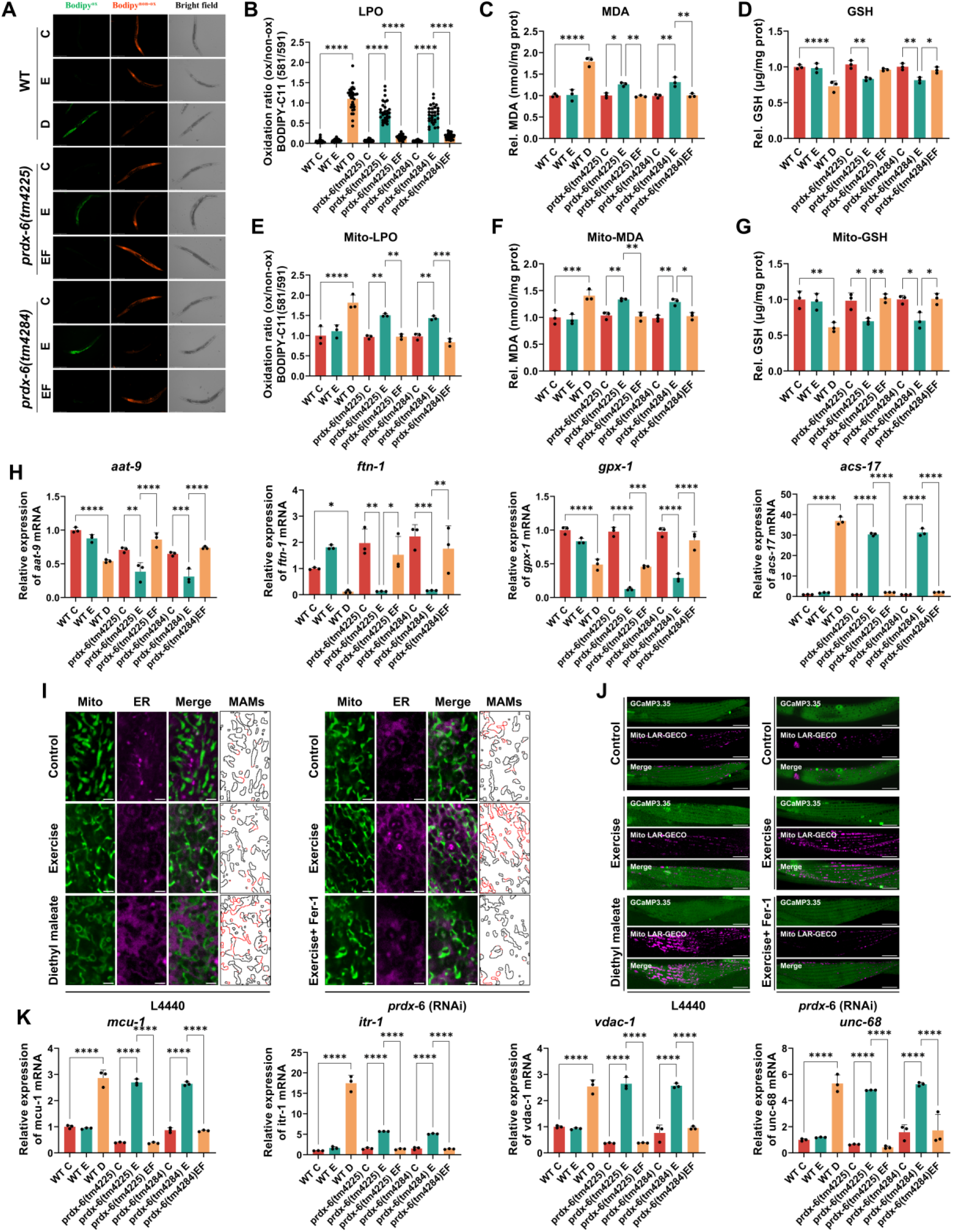
Exercise results in increased mitochondrial lipid peroxidation, increased MAMs assembly and mitochondrial Ca^2+^ levels in *prdx-6* mutants. Measurements were conducted on day 6 adult nematodes following the indicated treatments: controls (C), exercise (E), diethyl maleate (D) or exercise combined with ferrostatin-1 (EF). Panels A-H and K were performed in Wild-type and *prdx-6* mutant strains (*tm4225* and *tm4284*); panels I and J were performed in *vkEx2674[pnhx-2CemOrange2::PISY-1; pmyo-2GFP]; zcIs17[pges-1mitGFP]* MAM reporter and *goeIs3[pmyo-3SL1::GCamP3.35::SL2; unc-119(+)];aceIs1[pmyo-3mitoLAR-GECO; pmyo-2::RFP]* calcium reporter treated with either empty vector (L4440) or *prdx-6* (RNAi). **(A, B)** Lipid peroxidation by BODIPY-C11 (581/591). A, representative images showing the oxidised (green) and non-oxidised (red) signals with brightfield images. B, quantification of the whole animal oxidised to non-oxidised ratio. Scale bars, 275 µm. **(C, D)** Whole animal MDA and GSH. (**E)** Mitochondrial lipid peroxidation by BODIPY-C11, expressed as the ratio of oxidised ratio. **(F, G)** Mitochondrial MDA and GSH. **(H)** Relative mRNA expression levels of the ferroptosis genes *aat-9*, *ftn-1*, *gpx-1* and *acs-17*. **(I)** Representative images of MAMS using the *vkEx2674[pnhx-2CemOrange2::PISY-1; pmyo-2GFP]; zcIs17[pges-1mitGFP]* reporter under control (L4440) and *prdx-6* RNAi conditions, showing mitochondria, ER, the merge and the contact site mask. Scale bars, 2 µm. **(J)** Representative images of cytoplasmic and mitochondrial calcium using *goeIs3[pmyo-3SL1::GcamP3.35::SL2; unc-119(+)]; aceIs1[pmyo-3mitoLAR-GECO; pmyo-2::RFP]* under control (L4440) and *prdx-6* RNAi conditions. Scale bars, 20 µm. **(K)** Relative mRNA expression levels of the calcium channel genes *mcu-1*, *itr-1*, *vdac-1* and *unc-68*. Data presented as mean ± SEM. For B, n = 30; for C-H and K, n = 3. Significance was determined by one-way ANOVA. * *p* < 0.05, ** *p* < 0.01, *** *p* < 0.001, **** *p* < 0.0001.

### Ferroptosis phenotype in *C. elegans prdx-6* mutants following exercise results in mtDNA release and inflammatory signalling

As demonstrated in the myoblast model, knockdown of PRDX6 or inhibition of its phospholipase A2 activity resulted in mtDNA release and activation of inflammatory signalling. Cytoplasmic and mitochondrial fractions of the different strains following exercise or DEM treatments were prepared and the purity of each fraction was confirmed (Fig. 8A). qPCR analysis of mitochondrial DNA encoded genes (*cytb*, *cox-1* and *nd1*) in cytoplasmic fractions was performed. The results highlight that DEM in the wild type strain and exercise in the *prdx-6* mutants promoted the release of mtDNA into cytoplasmic fractions (Fig. 8B-D). In *C. elegans* the p38 MAPK pathway is activated in response to elevated stress [47]. Western blotting for phospo-p38 MAPK revealed increased levels in the wild type strain following DEM treatment and following exercise in the *prdx-6* mutant strains which could be alleviated with Fer-1 (Fig. 8E). Moreover, this was accompanied by increased expression levels of genes involved in the inflammatory response *sysm-1*, *irg-4*, *irg-5*, *K08D8.4* and *K08D8.5* [48–50] (Fig. 8F-J). Reporter strains for innate immune response in combination with RNAi knockdown of *prdx-6* were also analysed. Consistent with the previous results, treatment of wild type worms with DEM increased activation of reporter strains *irg-4p*::GFP, *irg-5p*::GFP and *sysm-1p*::GFP, similar to the exercise intervention with knockdown of *prdx-6*, which could be rescued with Fer-1 (Fig. 8K). In summary and similar to the cell model, loss of PRDX-6 and following a physiological stress results in the release of mtDNA and inflammatory signalling.

**Fig. 8.**
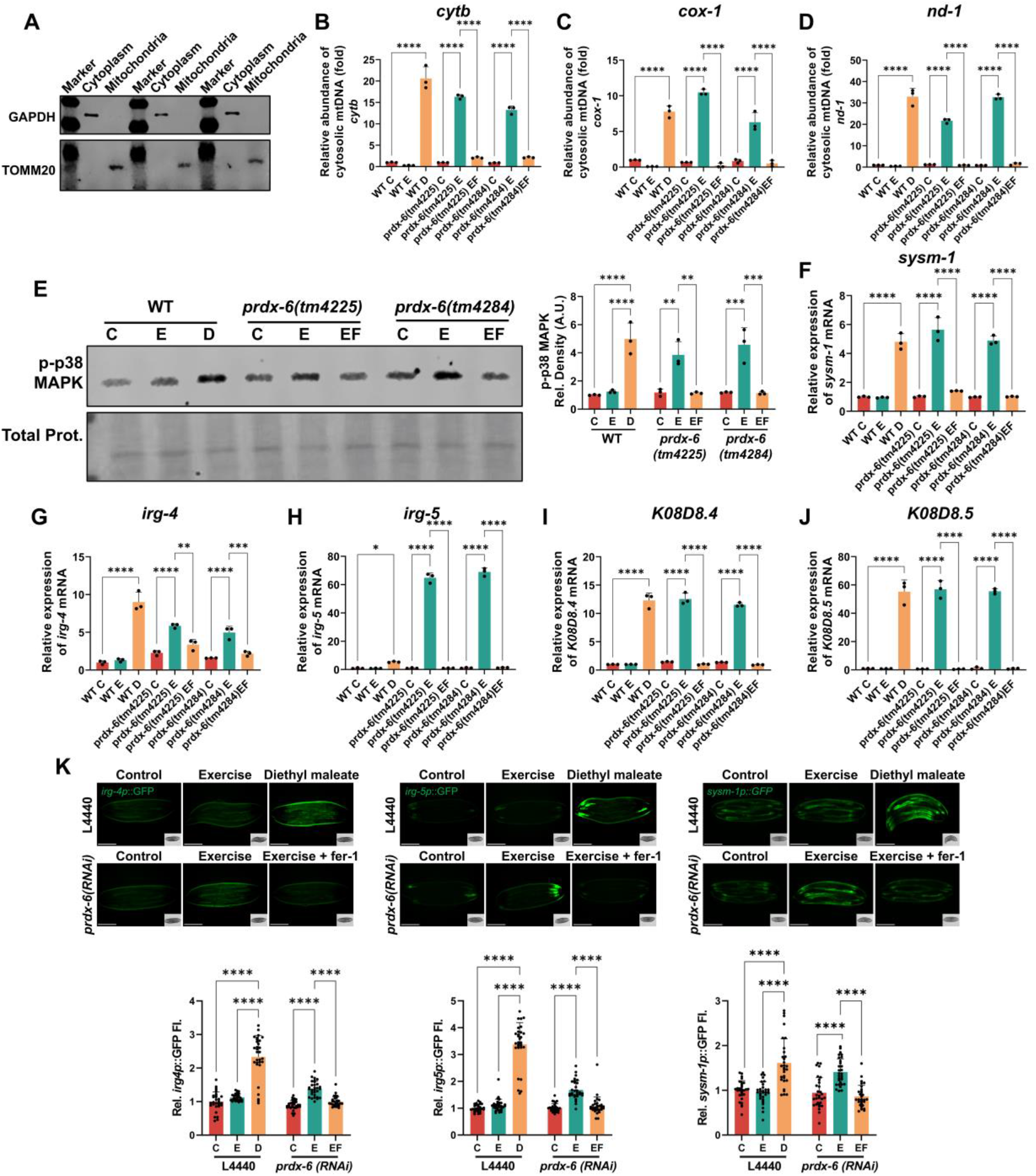
Exercise intervention in *prdx-6* mutant *C. elegans* results in mtDNA release and inflammatory signalling. Measurements were conducted on day 6 adult nematodes following the indicated treatments: controls (C), exercise (E), diethyl maleate (D) or exercise combined with ferrostatin-1 (EF). Panels A-J were performed in wild-type and *prdx-6* mutant strains (*tm4225* and *tm4284*); panel K used *prdx-6* RNAi with L4440 as the empty vector control. **(A)** Immunoblots confirming the purity of cytoplasmic and mitochondrial fractions, using GAPDH (cytoplasm) and TOMM20 (mitochondria) as markers. **(B-D)** qPCR quantification of the mtDNA encoded genes *cytb*, *cox-1* and *nd-1* in cytoplasmic fractions. **(E)** Immunoblot and quantification of phosphorylated p38 MAPK, with total protein as the loading control. **(F-J)** Relative mRNA expression levels of the innate immune genes *sysm-1*, *irg-4*, *irg-5*, *K08D8.4* and *K08D8.5*. **(K)** Innate immune reporter activation in animals treated with empty L4440 or *prdx-6* RNAi. Representative images and quantification of relative GFP fluorescence for the *irg-4p*::GFP, *irg-5p*::GFP and *sysm-1p*::GFP reporters under the conditions indicated. Scale bars, 275 µm. Data presented as mean ± SEM. For B-J n = 3; for K, n = 30. Significance was determined by one-way ANOVA. * *p* < 0.05, ** *p* < 0.01, *** *p* < 0.001, **** *p* < 0.0001.

### Prevention of excessive mitochondrial Ca^2+^ can protect cells from ferroptosis induced inflammatory signalling

In both the myoblast cell model and *C. elegans* strains both ferroptosis induction in control groups or a physiological stress in models lacking Peroxiredoxin 6 resulted in increased MERCS assembly that was associated with increased mitochondrial Ca^2+^ levels, mtDNA release and inflammatory signalling. In order to determine if inhibiting the elevated mitochondrial Ca^2+^ levels could rescue this phenotype, RNAi knockdown of the mitochondrial calcium uniporter (*mcu-1*) in the *prdx-6* mutant strains was performed. Similar to inclusion of the ferroptosis inhibitor Fer-1, RNAi knockdown of *mcu-1* rescued the decreased longevity following exercise in both *prdx-6* mutant strains (Fig. 9A). Knockdown of *mcu-1* also rescued the increased phospo-p38 MAPK following exercise in the *prdx-6* mutant strains (Fig. 9B). Furthermore, the RNAi mediated knockdown of *mcu-1* also prevented the exercise induced cytoplasmic release of mtDNA in the *prdx-6* mutant strains (Fig. 9C-E), the inflammatory signalling response as assessed by qPCR of inflammatory genes (Fig. 9F-J) and inflammatory reporter strains (Fig. 9K).

**Fig. 9.**
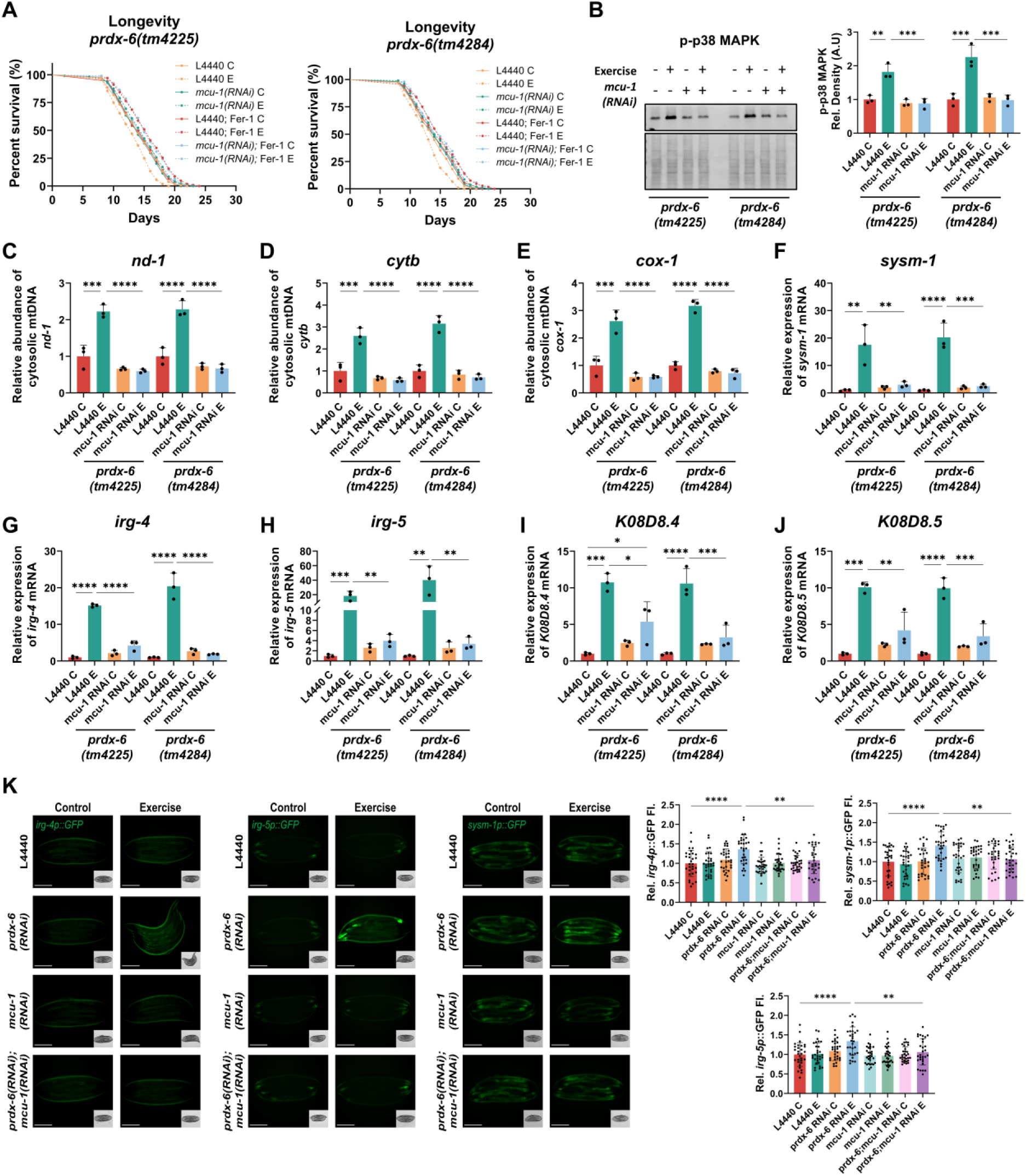
Knockdown of *mcu-1* rescues the exercise-induced inflammatory phenotype in *prdx-6* mutant *C. elegans*. Panels A-J, *prdx-6* mutant strains *tm4225* and *tm4284* are empty L4440 or *mcu-1* RNAi treated: controls (C), exercise (E), ferrostatin-1 (Fer-1). Panel K used single and combined RNAi knockdown of *prdx-6* and *mcu-1*. **(A)** Kaplan-Meier survival curves for lifespan of *prdx-6*(*tm4225*) and *prdx-6*(*tm4284*). **(B)** Immunoblot and quantification of phosphorylated p38 MAPK in each strain fed control or *mcu-1* RNAi, with total protein as the loading control. **(C-E)** qPCR quantification of the mtDNA encoded genes *nd-1*, *cytb* and *cox-1* in cytoplasmic fractions. **(F-J)** qPCR analysis of the innate immune genes *sysm-1*, *irg-4*, *irg-5*, *K08D8.4* and *K08D8.5*. **(K)** Innate immune reporter activation. Representative images and quantification of relative GFP fluorescence for the *irg-4p*::GFP, *irg-5p*::GFP and *sysm-1p*::GFP reporters in Wild-type animals fed control (L4440), *prdx-6* RNAi, *mcu-1* RNAi or combined *prdx-6* and *mcu-1* RNAi, following the 5-day swimming exercise. Scale bar*s,* 275 µm. Data presented as mean ± SEM. For A, n = 105; and *p* values were determined by the log-rank (Mantel-Cox) test. For B-J, n = 3; for K, n = 30. Significance for B-K was determined by two-way ANOVA. * *p* < 0.05, ** *p* < 0.01, *** *p* < 0.001, **** *p* < 0.0001.

In summary using both a cell myoblast model treated with an acute and relatively low concentration of H_2_O_2_ and the whole organism *C. elegans* subjected to a physiological exercise stress that promotes beneficial adaptive responses, we have identified the crucial role of Peroxiredoxin 6 in preventing ferroptosis induced signalling. In both models lacking Peroxiredoxin 6, following a relatively mild physiologically relevant stress there was increased mitochondrial fragmentation, increased MERCS assembly followed by increased mitochondrial Ca^2+^. Ultimately this excess mitochondrial Ca^2+^ promoted mtDNA release into the cytoplasm and the initiation of an inflammatory signalling response. Moreover, preventing the excessive mitochondrial Ca^2+^ levels could rescue this phenotype. Together the data highlights that under stress conditions Peroxiredoxin 6 localises to mitochondria and plays an essential role in the prevention of lipid peroxidation and ferroptosis signalling.

## Discussion

PRDX6 is a multifunctional enzyme with a range of distinct catalytic activities including repair of peroxidised cell membranes and regulation of lipid homeostasis [21, 25]. The role of PRDX6 in regulating the repair of lipid hydroperoxides and influencing sensitivity to ferroptosis has previously been identified [20, 51]. In this study using both a cell model and the whole organism *C. elegans*, it was demonstrated that PRDX6 localises to mitochondria under conditions of physiological stress and prevents lipid peroxidation and the initiation of ferroptosis signalling. In the absence of PRDX6 there was increased mitochondrial lipid peroxidation, that promoted increased MERCS formation and elevated mitochondrial Ca^2+^ levels, ultimately this resulted in mtDNA release and inflammatory signalling. The results highlight a key role of PRDX6 in preventing mitochondrial lipid peroxidation and inflammatory signalling under adaptive stress conditions.

The role of PRDX6 in protecting against ferroptosis induced cell death has been described in a cancer models and skeletal muscle [19–21, 28]. Both PRDX6 and GPX4 were suggested as the only two enzymes able to reduce phospholipid hydroperoxides at a significant rate [18, 19]. The levels of both these proteins have been intricately linked in the repair of oxidised membranes, the reduction of hydroxyl precursors by GPX4 are hydrolysed by the phospholipase activity of PRDX6 reducing the toxicity of oxidised phospholipids [19]. Following ferroptosis induction with erastin or GSH depletion using BSO, PRDX6 can modulate GPX4 localisation and function, as PRDX6 Cys47 can form an intramolecular disulphide with GPX4 [19]. In this study, under conditions of reduced levels of PRDX6, no changes in GPX4 levels were observed in control conditions, however following H_2_O_2_ or erastin treatment the levels of GPX4 significantly decreased, suggesting that under stress conditions PRDX6 is required for the stability of GPX4. Targeting or the modification of PRDX6 activities has received increasing attention, as an increase in PRDX6 levels has been described in a number of different cancer tissues which was associated with increased resistance to chemotherapy induced cell death [19, 51]. However, the changes in PRDX6 levels are tissue dependent, for example in skeletal muscle mitochondrial localised PRDX6 levels have been found to decrease with age [28]. Loss of mitochondrial function is one of the hallmarks of ageing, and in skeletal muscle the loss of mitochondrial function underlies the loss of muscle mass and function [52]. Decreased mitochondrial localised PRDX6 with age is associated with increased mitochondrial lipid peroxidation and decreased mitochondrial function [28]. Site directed mutation of PRDX6 phospholipase activity, results in an impaired ability to repair oxidised lipids and loss of mitochondrial bioenergetics highlighting its key enzymatic function [28]. Both the peroxidase and phospholipid hydroperoxidase activities of PRDX6 are required to alleviate lipid peroxidation [28]. Studies of PRDX6 knockdown using cell models combined with lipidomics, reported an increase in sphingomyelins and acylcarnitines resulting in increased PUFAs and lipid droplets [21]. Experimental approaches in rodents have confirmed the phospholipase activity of PRDX6 is required to maintain mitochondrial function and lipid homeostasis [28].

It has also been described that PRDX6 can act as a Selenium binding protein, facilitating the incorporation of selenocysteine into the catalytic site of GPX4 and regulation of its activity [20, 51]. A number of studies in cancer models have reported that loss of PRDX6 increases ferroptosis sensitivity due to the inhibition of selenoprotein synthesis required for GPX4, while other studies have reported no reduction in GPX4 levels [20, 28, 51]. In *C. elegans*, the levels of *gpx-1* decreased in the wild type strain following glutathione depletion with DEM and following exercise in the *prdx-6* mutants strains which could be rescued with the ferroptosis inhibitor, Fer-1. Unlike mammalian species, there is only one identified selenoprotein in *C. elegans*, Thioredoxin reductase-1, which suggests that the role of PRDX6 in protection against ferroptosis is not due to its role in selenium binding in these models.

The link between MERCS and lipid peroxidation has recently been identified, these areas are highly enriched sites for lipid peroxidation during the induction of ferroptosis [15]. MERCS are highly dynamic structures and key regulators of intracellular homeostasis by mediating inter-organelle crosstalk and the exchange of lipids, Ca^2+^, metabolites and iron, in response to biological perturbations [10]. MERCS are also key determinants of mitochondrial dynamics as they mark the sites of mitochondrial fusion and fission [7, 9]. Disruption of mitochondrial dynamics has been reported in a wide variety of age-related diseases including neurodegeneration and sarcopenia, associated with an accumulation of dysfunctional mitochondria [53]. An increase in MERCS formation has been proposed to be involved in cell senescence and in models of neurodegenerative disease [13, 54]. In skeletal muscle, sarcomeres are surrounded by mitochondria and the sarco/endoplasmic reticulum, essential for Ca^2+^ regulation, however decreased MERCS formation has been reported with age [55], indicating that MERCS assembly is context dependent. This context dependence is reinforced by our previous demonstration that adaptive ER stress promotes PERK-dependent MERCS assembly that supports mitochondrial remodelling and extends lifespan [6]. Taken together with the present findings, this argues that the outcome of contact-site remodelling is determined not by the abundance of MERCS alone but by their qualitative state, including the burden of peroxidised phospholipids and the magnitude of the Ca^2+^ flux they support. The change in cellular architecture from development to ageing could inhibit the dynamics of MERCS turnover. Energetic stress and subsequent AMPK activation have also been demonstrated in cell models to promote autophagy and MERCS formation [56]. Previously it was demonstrated that an exercise protocol in *C. elegans* generates energetic stress, induces mitochondrial remodelling that ultimately results in an adaptive response and improved bioenergetic profile [31]. However, following ferroptosis the contact sites between the ER and mitochondria expand ultimately resulting in an acute increase in mitochondrial ROS generation and increased mitochondrial fission [15]. In both the cell and *C. elegans* models lacking PRDX6 a similar increase in MERCS assembly was identified following an adaptive physiological stress that was accompanied by increased lipid peroxidation and mitochondrial fragmentation. The increased lipid peroxidation and ferroptosis signalling following MERCS expansion described here is also supported by work which has identified MERCS as key hubs in the transport of iron into mitochondria [10]. Furthermore, MERCS expansion results in increased levels of mitochondrial Ca^2+^, which has been demonstrated to result in opening of the permeability transition pore (PTP) [57]. The PTP is an essential regulator of mitochondrial redox homeostasis, as sustained opening on the PTP can increase mtROS and promote the activation of cell death pathways [57]. Although, this study did not determine altered PTP opening, the increase in mitochondrial Ca^2+^ was associated with increased release of mtDNA when PRDX6 was absent under physiological stress conditions. Inflammation in response to harmful stimuli occurs in response to the cellular recognition of DAMPs, one of which is the release of mtDNA due to its proteobacterial origin [58]. The results presented here confirmed a large increase in the innate immune response following mtDNA release in both the cell and *C. elegans* models. Interestingly, the inhibition of mitochondrial Ca^2+^ uptake in the *C. elegans* model by RNAi knockdown of *mcu-1*, prevented the release of mtDNA and subsequent inflammatory signalling that ultimately rescued the reduced longevity of strains lacking *prdx-6* following the exercise intervention. Knockdown of *mcu-1* or pharmacological inhibition of the MCU complex in early adulthood has been demonstrated to increase longevity and activity of old worms, although can impair survival in early adulthood [59]. Together, these results highlight the context dependency of regulating mitochondrial Ca^2+^ levels, a number of TCA enzymes are Ca^2+^ dependent but excess mitochondrial Ca^2+^ generates increased ROS and opening of the PTP resulting in activation of cell death pathways [60, 61].

The results presented identify a crucial role of PRDX6 in preventing mitochondrial lipid peroxidation, increased MERCS assembly and activation of ferroptosis under physiological stress conditions. However, there are a number of limitations in the current study, in particular identifying the distinction between adaptive and maladaptive MERCS assembly. Not only the abundance of MERCS is important but also the distance between the organelles, tight junctions (<10 nm) have been suggested to promote lipid signalling while wider junctions (∼ 30-50 nm) promote Ca^2+^ transfer [9, 61]. Electron volume microscopy and correlative imaging would resolve the width of the gap between the membranes and allow the wider contacts that favour Ca^2+^ exchange to be separated from the narrower contacts that favour lipid transfer, and these structural measures could be combined with functional readouts of the Ca^2+^ and lipid flux that pass through the contacts sites. Also, the molecular basis of the translocation of PRDX6 to mitochondria has not been resolved, since PRDX6 lacks a canonical mitochondrial targeting sequence and the trafficking machinery responsible for its recruitment remains to be defined.

In summary, the results presented in this study identify PRDX6 as a key enzyme in regulating mitochondrial lipid peroxidation and prevention of ferroptosis in response to physiological stress. The loss of PRDX6 or inhibition of its phospholipase activity results in increased mitochondrial MERCS assembly and mitochondrial Ca^2+^ levels, that result in increased mitochondrial fragmentation and mtDNA release ultimately leading to an inflammatory response. The results highlight that modulation of PRDX6 activities has important translational applications in conditions such as cancer that can sensitise resistant tumours to cell death but also in age-related conditions associated with increased mitochondrial dysfunction and inflammation.

## Methods

### Cell culture and treatments

C2C12 myoblasts were maintained in growth medium (GM) comprising high-glucose (4.5 g/L) DMEM supplemented with 10% fetal bovine serum (FBS) and 1% penicillin-streptomycin (P/S). For myogenic differentiation, cells were switched to differentiation medium (DM; DMEM with 2% horse serum and 1% P/S) for 7 days. To test the effects of H₂O₂ on myogenesis, sub confluent myoblasts were exposed to graded H₂O₂ (0, 5, 10, 25, 50 and 100 µM) for 10 min, followed by replacement with fresh GM and a 24 h recovery period before induction of differentiation. Based on the dose-response experiments, 25 μM H₂O₂ for 10 min was selected for subsequent experiments. To inhibit the calcium-independent phospholipase A₂ (aiPLA₂) activity of PRDX6, cells received 10 µM MJ33 from 1 h before H₂O₂ exposure, with the inhibitor maintained throughout treatment and the 24 h recovery to ensure sustained inhibition. Ferroptosis lipid peroxidation was induced with 1 µM erastin for 4 h, after which cells were returned to fresh GM and allowed to recover for 24. The lipophilic radical-trapping antioxidant ferrostatin-1 (Fer-1, 10 µM) was applied during H₂O₂ or erastin exposure and sustained through the 24 h recovery to block lipid peroxidation.

### Cell transfection

For transient gene silencing, proliferating cells were transfected with siRNA using Lipofectamine 2000 in serum-free DMEM for 5 h, with a final siRNA concentration of 50 nM. Lipofectamine 2000-only treatment was used as the transfection control. After transfection, the medium was replaced with GM, and cells were allowed to recover overnight before downstream treatments. For plasmid transfection, cells were transfected with Cox8-EGFP-mCherry or ER-mit splitFAST plasmids using Lipofectamine 2000 to assess mitophagy and mitochondria-endoplasmic reticulum contact site formation, respectively [34, 62]. Briefly, 500ng plasmid DNA was complexed with Lipofectamine 2000 in serum-free DMEM for 20 min at room temperature and added to the cells for 5 h, after which the medium was replaced with GM and cells were allowed to recover overnight before subsequent experiments.

### *C. elegans* maintenance and treatments

*C. elegans* strains were maintained at 20°C in the dark on standard nematode growth medium (NGM) plates seeded with *Escherichia coli* OP50 as food source. The strains used in this study are listed in Supplementary Table 1. Synchronized worm populations were obtained using standard bleaching. To induce lipid peroxidation, worms were transferred to OP50-seeded NGM plates containing 5 mM diethyl maleate (DEM) in the agar and maintained at 20°C for 16 h. For rescue experiments, worms were treated with agar containing ferrostatin-1 (Fer-1; 200 μM), either alone or in combination with DEM, under the same conditions. After treatment, animals were collected for downstream analyses.

### Western blotting

Total protein was extracted from C2C12 cells and *C. elegans* samples using RIPA lysis buffer. When the redox state of peroxiredoxins was examined, samples were lysed using an alkylating lysis buffer (150 mM NaCl, 20 mM Tris-HCl pH 7.5, 1 mM EDTA pH 8.3, 0.5% SDS, 1% Triton X-100, and 100 mM N-ethylmaleimide). Protein extracts were quantified using the Bradford assay. Western blotting was performed using equal amounts of protein separated on reducing SDS-PAGE gels. Proteins were transferred to membranes using a semi-dry transfer system, total protein loading was assessed by Ponceau S staining. Membranes were washed with TBS-T, blocked for 1 h at room temperature with 5% non-fat milk in TBS-T, or with 5% BSA in TBS-T when phosphorylated proteins were detected, and then incubated overnight at 4°C with primary antibodies diluted 1:1000 in the appropriate blocking buffer. After washing, membranes were incubated with fluorescent secondary antibodies diluted 1:15,000 in TBS-T for 1 h at room temperature protected from light. Signals were acquired using an Odyssey Fc imaging system and quantified using ImageJ. Target protein abundance was normalized to total protein loading based on Ponceau S staining. Uncropped Western blots are showed in the Supplementary data file.

### Quantitative real-time PCR (qRT-PCR)

Total RNA was isolated from C2C12 cells and *C. elegans* using a standard TRIzol-based extraction method, and RNA concentration and purity were assessed using a NanoDrop 2000 spectrophotometer. For cDNA synthesis, 500 ng total RNA was diluted with nuclease-free water to 10 μL, mixed with 1 μL random hexamers, heated at 65°C for 10 min, and immediately chilled on ice. Reverse transcription was then performed by adding 9 μL of master mix containing 4 μL reverse transcription buffer, 2 μL DTT, 1 μL dNTP mix, 1 μL SuperScript II reverse transcriptase, and 1 μL RiboLock RNase inhibitor. Reactions were incubated at 42°C for 60 min, after which cDNA was diluted 10-fold with 180 μL nuclease-free water for qRT-PCR analysis. qRT-PCR was performed using Fast SYBR Green in a final volume of 5.5 μL containing 2.5 μL 2× Fast SYBR Green qPCR Master Mix, 0.25 μL each primer, and 2.5 μL diluted cDNA template. Amplification was conducted under the following conditions: 95°C for 1 min, followed by 40 cycles of 95°C for 3 s and 60°C for 30 s. All reactions were performed in triplicate. Relative gene expression was calculated using the 2-ΔΔCt method, with *B2M* used as the internal reference gene for C2C12 samples and *cdc-42* used for *C. elegans* samples. The sequences of primers used in this study are listed in Supplementary Table 1.

### Mitochondrial and Cytosolic fractionation isolation

Mitochondrial and cytosolic fractions were isolated from cultured cells and *C. elegans* by differential centrifugation [58]. For cultured cells, samples were harvested, washed with ice-cold PBS, and resuspended in 200 μL freshly prepared digitonin buffer containing 50 mM HEPES pH 7.4, 150 mM NaCl, 20 μg/mL digitonin, and 1% protease inhibitor cocktail. Cells were incubated on a tube rotator for 10 min at 4°C to permeabilize the plasma membrane, followed by 10-15 strokes using a pre-chilled tight-fit Dounce homogenizer. For *C. elegans*, worms were collected, washed three times with M9 buffer and once with ice-cold PBS, and resuspended in 3 mL ice-cold mitochondrial isolation buffer containing 50 mM KCl, 70 mM sucrose, 0.1 mM EDTA pH 8.0, 5 mM Tris-HCl pH 7.4, and 1% protease inhibitor cocktail. Worm suspensions were homogenized on ice using 21 gentle Dounce strokes. Cell and worm homogenates were then clarified by centrifugation at 800 × g for 10 min at 4°C to remove unbroken cells, nuclei, and large debris. To improve mitochondrial recovery, the pellet was re-homogenized and centrifuged at 1,200 × g for 3 min at 4°C, and the resulting supernatant was further centrifuged at 1,200 × g for 3 min to remove residual heavy contaminants. Cleared supernatants were pooled and centrifuged at 11,000 × g for 15 min at 4°C to obtain the mitochondrial pellet. The remaining supernatant was further centrifuged at 21,000 × g for 30 min at 4°C, and the final supernatant was collected as the cytosolic fraction. The mitochondrial pellet was washed three times with ice-cold isolation buffer before downstream analysis. Fraction purity was routinely verified by western blotting using TOM20 and GAPDH as mitochondrial and cytosolic markers, respectively.

### Cytosolic mtDNA measurement

Cytosolic mtDNA was measured to assess mitochondrial DNA release into the cytosol. Total DNA was extracted in parallel from whole-cell lysates and cytosolic fractions using a standard protocol adapted from [8]. For cultured cells, whole-cell lysates were prepared by directly resuspending cell pellets in SDS lysis buffer containing 20 mM Tris pH 8.0, 1% SDS, and 1% protease inhibitor cocktail, followed by heating at 95°C for 15 min. For *C. elegans*, worm pellets were resuspended in the same SDS lysis buffer, subjected to three freeze–thaw cycles, vigorously vortexed, and heated at 95°C for 15 min. Cytosolic fractions from cells and *C. elegans* were prepared as described above. For DNA extraction, 200 μL of each whole-cell or cytosolic fraction was treated with 1 μL RNase A (10 mg/mL) at 37°C for 1.5 h, followed by digestion with 2 μL proteinase K (20 mg/mL) at 55°C for 1 h. Samples were then extracted twice with an equal volume of chloroform/isoamyl alcohol (24:1, v/v), with vigorous vortexing for 1 min and centrifugation at 21,000 × g for 5 min at room temperature after each extraction. The aqueous phase was recovered and DNA was precipitated by adding one-tenth volume of 7.5 M ammonium acetate, 1 μL glycogen (20 μg), and 2.5 volumes of 100% ethanol, followed by incubation at −80°C for 1 h. DNA was pelleted by centrifugation at maximum speed for 20 min at 4°C, washed twice with 300 μL 95% ethanol, air-dried for 5 min, and resuspended in 50 μL nuclease-free water. DNA concentration and purity were determined using a NanoDrop 2000 spectrophotometer. The relative abundance of mtDNA encoded genes in the cytosol was quantified by qRT-PCR using primers targeting representative mitochondrial loci, such as *mt-Cytb*, and normalized to nuclear DNA loci (*Tert* for samples from cells, *tbb-2* for samples from *C. elegans*), as described previously [58].

### MDA quantification

MDA content was determined using a commercial MDA Content Assay Kit (BC0025, Solarbio, Beijing, China) according to the manufacturer’s instructions. Briefly, the collected cell, *C. elegans*, or mitochondrial samples were homogenized or lysed in the extraction buffer provided with the kit on ice and centrifuged at 8,000 × g for 10 min at 4°C. The resulting supernatant was mixed with the MDA detection working solution, followed by incubation in a boiling water bath at 100°C for 60 min. After rapid cooling on ice, the reaction mixture was centrifuged at 12,000 × g for 10 min, and the supernatant was transferred to a 96-well plate. Absorbance was measured at 532 and 600 nm using the Hidex plate reader. MDA levels were calculated from the corrected absorbance difference (A532 - A600) according to the kit formula and normalized to the corresponding protein concentration.

### GSH quantification

GSH content was measured using a commercial GSH Content Assay Kit (BC1175, Solarbio, Beijing, China) according to the manufacturer’s instructions. Briefly, the collected cell, *C. elegans*, or mitochondrial samples were homogenized in the extraction reagent provided with the kit under ice-cold conditions and centrifuged at 12,000 × g for 10 min at 4°C. The resulting supernatant was collected for analysis. For the assay, 20 μL of sample supernatant, GSH standard, or distilled water blank was mixed with 140 μL of reagent II and 40 μL of reagent III in a 96-well plate. After incubation at room temperature for 2 min, absorbance was measured at 412 nm using the Hidex plate reader. GSH concentration was calculated from a standard curve generated with serially diluted GSH standards and normalized to cell number or protein concentration.

### Immunocytochemistry for myosin heavy chain (MF20)

MF20 immunostaining was performed to assess myogenic differentiation. After 7 days of differentiation, C2C12 cells were rinsed with PBS, fixed in ice-cold 100% methanol for 5 min, and blocked with PBS containing 10% horse serum for 1 h at room temperature. Cells were incubated overnight at 4°C with MF20 antibody diluted 1:1000 in PBS containing 2% horse serum, followed by Alexa Fluor 488-conjugated anti-mouse secondary antibody diluted 1:2000 for 45 min at room temperature in the dark. Nuclei were counterstained with DAPI diluted 1:10,000 for 5 min. Images were acquired using an EVOS M7000 imaging system at 10× magnification, with at least five non-overlapping fields captured per well from three independent biological replicates. Myotube diameter, MF20-positive area fraction, and fusion index were quantified using ImageJ.

### Live-cell mitochondrial staining

Live-cell fluorescence staining was used to assess mitochondrial mass, mitochondrial reactive oxygen species production, and mitochondrial membrane potential. After treatments, cells were washed with pre-warmed PBS and stained with Hoechst 33342 diluted 1:10,000 for 5 min at 37°C. Mitochondrial mass was evaluated using MitoTracker Green at 200 nM for 30 min at 37°C. Mitochondrial ROS and membrane potential were assessed using MitoSOX Red at 5 μM for 15 min and TMRM at 100 nM for 15 min, respectively. After staining, cells were washed thoroughly with PBS and imaged live using an EVOS M7000 fluorescence microscope at 40× magnification. Fluorescence intensity was quantified using ImageJ.

### Calcium imaging

Mitochondrial calcium was examined using Rhod-2 AM together with MitoTracker Green, the latter used to delineate the mitochondrial compartment occupied by the Rhod-2 signal. Cells were washed in pre-warmed PBS and co-loaded with 5 µM Rhod-2 AM and 200 nM MitoTracker Green in serum-free DMEM for 45 min at 37 °C, after which the loading solution was replaced with GM for a further 30 min at 37 °C to allow de-esterification. Cytosolic calcium was measured using Fluo-4 AM. Thapsigargin (1 µM, 30 min, 37 °C), an inhibitor of the sarcoplasmic/endoplasmic reticulum Ca^2+^-ATPase that mobilises ER calcium stores, was applied as a positive control [6]. Cells in all conditions were then loaded with 3 µM Fluo-4 AM for 45 min at 37 °C in the dark, switched to GM for 30 min, washed in PBS to remove extracellular dye and imaged immediately on the EVOS M7000.

### BODIPY 581/591 C11 staining

Lipid peroxidation was assessed with BODIPY 581/591 C11, with oxidation reported as the green-to-red emission ratio [7]. For whole-cell measurements, treated cells were loaded with 1 µM BODIPY C11 in pre-warmed DPBS for 30 min at 37 °C in the dark, washed and imaged immediately on an EVOS M7000 at 60× magnification. Reduced and oxidised BODIPY signals were acquired in the red and green channels, respectively, and lipid peroxidation was quantified in ImageJ as the green-to-red fluorescence ratio after background correction. For mitochondrial lipid peroxidation, cells were co-stained with 1 µM BODIPY C11 and 200 nM MitoTracker Deep Red. In ImageJ, the oxidised-BODIPY (green) and mitochondrial (far-red) channels were thresholded into binary masks using a single strategy applied across all images within an experiment. The two masks were intersected with the Image Calculator “AND” operation to define lipid-peroxidised mitochondrial regions. The percentage of lipid-peroxidised mitochondria was calculated as the BODIPY C11-positive mitochondrial area divided by the total mitochondrial area. The overlap mask was inverted to display peroxidised mitochondria as black structures on a white background for visualisation.

### ER-mitochondria contact site imaging (ER-mito splitFAST)

Mitochondria-ER contact sites (MERCS) were visualised by transfecting the ER-mito splitFAST reporter [34], with mitochondria counterstained using MitoTracker Deep Red. Transfection and mitochondrial staining were performed as described above, and images were acquired on an EVOS M7000 at 60× magnification. In ImageJ, the MitoTracker and splitFAST channels were background-subtracted by the rolling-ball method, smoothed with a Gaussian filter (sigma 1 pixel) and thresholded into independent binary masks using identical parameters within each experiment [34]. To quantify ER-mitochondria contact sites associated with mitochondria, the mitochondrial mask and ER-mito splitFAST mask were intersected using the Image Calculator “AND” function in ImageJ. The overlap area and total mitochondrial area were measured within individual cell regions of interest. The percentage of mitochondrial area associated with MERCS was determined as the overlap area divided by the total mitochondrial area.

### Immunofluorescence staining

For immunofluorescence staining, C2C12 myoblasts were cultured on laminin-coated coverslips. After treatments, live cells were stained with MitoTracker Deep Red at 200 nM for 30 min at 37°C in the dark, rinsed with PBS, and fixed with 4% paraformaldehyde for 15 min at room temperature. Cells were permeabilized with 0.25% Triton X-100 in PBS for 15 min, blocked with 5% BSA in PBS for 30 min, and incubated overnight at 4°C with primary antibodies diluted in 1% BSA according to the manufacturer’s instructions. After washing, cells were incubated with fluorophore-conjugated secondary antibodies diluted 1:2000 in PBS for 1 h at room temperature in the dark. Coverslips were mounted with Hydromount, left overnight at room temperature, and imaged using an EVOS M7000 fluorescence microscope. Images were analysed using ImageJ.

### Swimming exercise

Day 1 adult worms were washed three times with M9 buffer to remove bacteria, larvae, and debris. For the exercise group, cleaned worms were transferred to OP50-seeded NGM plates supplemented with 3 mL M9 buffer to allow swimming, whereas control worms were placed on standard OP50-seeded NGM plates without additional M9 buffer. Both groups were maintained at 20°C for 90 min per session. After each session, animals were collected with M9 buffer, allowed to settle by gravity, and transferred to fresh OP50-seeded NGM plates. The exercise intervention was performed twice daily, at approximately 09:00 and 15:00, for 5 consecutive days, after which worms were maintained at 20°C until the next intervention or downstream analysis [31].

### Lifespan assays

Lifespan assays were initiated following the final day of swimming exercise intervention. A total of 105 nematodes were randomly distributed across three seeded NGM plates supplemented with 10 μM 5-fluoro-2′-deoxyuridine (FUdR) to inhibit egg hatching. The number of live and dead nematodes was recorded every one to two days until all individuals died. Nematodes that crawled off the plates or burrowed into the agar were censored. Detailed survival data, including mean survival, and log-rank test *p* values, are summarised in Supplementary Table 2.

### Oxidative stress survival assays

Oxidative stress survival assays were performed on the day after the swimming exercise regimen. Paraquat and sodium arsenite were utilised to induce mitochondrial and cytoplasmic oxidative stress, respectively [63, 64]. Fresh aliquots of 250 μL of 100 mM paraquat or 5 mM sodium arsenite, diluted in M9 buffer, were dispensed into each well of a 24-well plate. Subsequently, 12 to 14 nematodes, totalling 50 nematodes per group, were randomly placed into each well. The number of surviving nematodes was recorded every one to two hours until all individuals died. Nematodes were considered dead if they failed to respond to gentle stimulation with a platinum wire picker. Detailed survival data, including mean survival, and log-rank test *p* values, are summarised in Supplementary Table 2.

### *C. elegans* swimming test (CeleST)

The CeleST assay was conducted one day after the swimming exercise regimen, as described previously [65]. Briefly, five nematodes were randomly selected and placed into a 40 μL droplet of M9 buffer, which were confined within a 10 mm circular barrier on a glass slide. Nematodes were allowed to settle in the M9 buffer for 20 s, then their movement was recorded for 30 seconds at 15 frames per second using a Nikon LV-TV microscope equipped with an OPTIKA C-P20CM camera at 1x magnification. Data processing and the determination of swimming metrics were performed using a custom MATLAB application. A minimum of 30 nematodes per condition were evaluated.

### RNA interference (RNAi)

RNA interference (RNAi) was performed using the bacterial feeding method. Briefly, *E. coli* HT115 strains carrying gene-specific dsRNA-expressing constructs were seeded onto NGM plates supplemented with IPTG (1 mM) and ampicillin (100 μg/mL) and incubated for 48 h at RT to induce dsRNA expression. Age-synchronized L1 larvae were placed on RNAi plates and maintained until adulthood, and their progeny were used for subsequent experiments. For double-gene knockdown, RNAi bacterial cultures were normalized to an optical density of 600 nm (OD600 = 50, stationary phase) and mixed at a 1:1 ratio before seeding. *E. coli* HT115 carrying the empty pL4440 vector was used as the negative control. Knockdown efficiency was assessed by qRT-PCR (Suppl. Fig. 5).

### C. elegans imaging

Unless otherwise indicated, *C. elegans* images were acquired using an EVOS M7000 imaging system. Quantification was performed using raw images without contrast enhancement in ImageJ. For live staining assays, worms were incubated with MitoSOX Red (10 μM, 1 h), or TMRE (200 nM, 6 h) to assess mitochondrial ROS, and mitochondrial membrane potential, respectively. After staining, worms were transferred to OP50-seeded NGM plates and allowed to feed for 2 h to clear unabsorbed dye from the gut. Animals were then immobilised on unseeded NGM plates using 40 mM levamisole and imaged at 10× magnification. All staining and imaging procedures were performed in the dark, and 30-45 worms per group were imaged.

To exam the lipid peroxidation in *C. elegans*, worms were collected and incubated with 10 µM BODIPY C11 in M9 buffer for 1 h at 20 °C in the dark with gentle agitation. Worms were returned to seeded NGM plates for 2 h to clear unabsorbed dye from the gut before mounting and imaging on the EVOS M7000. Lipid peroxidation was quantified in ImageJ after background subtraction. For isolated mitochondria, suspensions were normalised for protein content by Bradford assay, incubated with 5 µM BODIPY C11 for 30 min at RT in the dark and transferred to a black-walled, clear-bottom 96-well plate. Reduced (581/591 nm) and oxidised (488/510 nm) emissions were recorded on the Hidex plate reader, and the green-to-red ratio was normalised to mitochondrial protein content.

Mitochondrial content and morphology in body-wall muscle were assessed using the SJ4103: *zcIs14 [myo-3p::GFP(mit)]* reporter strain. For mitochondrial content, 30 worms per group were immobilized on unseeded NGM plates and imaged at 10× magnification. For mitochondrial morphology, worms were mounted on 4% agarose pads, and 130-150 images of body-wall muscle mitochondria between the pharynx and vulva were acquired at 60× magnification. Mitochondrial morphology was classified into five categories: class 1, abundant mitochondria forming a preserved tubular network; class 2, abundant mitochondria with network gaps and minor vesiculation; class 3, relatively sparse mitochondria with network gaps and increased vesiculation; class 4, sparse and disorganized mitochondria with minor vesiculation; and class 5, extremely sparse and disorganized mitochondria [66].

Mitophagy was assessed using the IR2539: *unc-119(ed3); Ex[myo-3p::tomm-20::Rosella; unc-119(+)]* strain. For each group, 45 worms were examined, and green and red fluorescence signals within the muscle-specific region between the pharynx and anterior intestine were acquired at 60× magnification. Mitophagy was quantified as the green-to-red fluorescence intensity ratio for each worm [31].

MAMs were analysed using the KPA427: *vkEx2674[pnhx-2CemOrange2::PISY-1; pmyo-2GFP]; zcIs17[pges-1mitGFP]* strain, which labels intestinal endoplasmic reticulum and mitochondria [46]. Dual-channel fluorescence images were acquired from the distal intestine using an Olympus FV3000 confocal microscope at 60× magnification. For each worm, two regions of interest measuring 10.45 × 10.45 μm were selected, and the mean value was used for statistical analysis. In ImageJ, ER and mitochondrial channels were converted into binary masks, filtered using a median filter with a radius of 0.5, and used to calculate the overlap between the ER mask and mitochondrial surface mask. MAMs were expressed as the percentage of mitochondrial surface area overlapping with the ER signal.

Calcium levels in body-wall muscle were assessed using genetically encoded calcium indicator strains. HBR4: *unc-119(ed3); goeIs3[pmyo-3::GCaMP3.35::unc-54 3′UTR; unc-119(+)]* was used to monitor overall calcium levels in body-wall muscle and imaged under the GFP channel at 10× magnification. ATU2301: *goeIs3[pmyo-3SL1::GCaMP3.35::SL2; unc-119(+)]; aceIs1[pmyo-3mitoLAR-GECO; pmyo-2::RFP]* was used to simultaneously assess cytosolic and mitochondrial calcium in body-wall muscle cells and imaged using an Olympus FV3000 confocal microscope at 60× magnification [46].

Innate immune pathway activation was examined using GFP transcriptional reporter strains, including AU306: *agIs44[irg-4p::GFP::unc-54 3′UTR; myo-2p::mCherry]*, AY101: *acIs101[irg-5p::GFP + rol-6(su1006)]*, and AU78: *agIs219[sysm-1p::GFP::unc-54 3′UTR + ttx-3p::GFP::unc-54 3′UTR] III*. These reporters were used to assess activation of intestinal immune effector genes regulated by the p38 PMK-1 pathway in *C. elegans* [50, 67].

## Statistical analysis

Statistical analyses were performed using GraphPad Prism version 10.5. The statistical test used for each experiment is indicated in the corresponding figure legend. Unless otherwise stated, data are presented as mean ± SEM from at least three independent biological replicates. For comparisons between two independent groups, data were analysed by Student’s t-test. For comparisons among more than two groups with a single independent variable or without a complete 2 × 2 factorial design, data were analysed by one-way ANOVA. For experiments with a complete 2 × 2 factorial design, two-way ANOVA was employed. For multiple treatment groups that did not constitute a complete factorial design, data were analysed by one-way ANOVA followed by Tukey’s multiple comparisons test. Survival curves from *C. elegans* lifespan and stress-resistance assays were generated using the Kaplan-Meier method and compared using the Mantel-Cox log-rank test. Pairwise survival comparisons were performed using log-rank tests with Holm correction for multiple testing. Categorical distributions were compared using the chi-square test. A *p*-value < 0.05 was considered statistically significant, with significance defined as \**p* ≤ 0.05, \*\**p* ≤ 0.01, \*\*\**p* ≤ 0.001, and \*\*\*\**p* ≤ 0.0001.

## Acknowledgements

We would like to sincerely thank Konstantonis Palikaras (University of Athens) for providing the KPA427 (MAM) reporter strain and ATU2301 Calcium reporter strain, Nektarios Tavernarakis (University of Crete) for providing the IR2539 mitophagy reporter strain and Read Pukkila-Worley (University of Massachusetts Chan Medical School) for providing the AU306 IRG-4 reporter strain. We would also like to sincerely thank Riccardo Filadi and Paola Pizzo (University of Padua) for sharing the splitFAST plasmids for analysis of MERCS and Corina Madreiter-Sokolowski (Medical University of Graz) for sharing the *mcu-1* RNAi. PL (202206370063), YZ (202406220057), JT (202306370005) and QX (202006370047) studentships are funded by the Chinese Scholarship Council (CSC), the JCCM studentship is funded by the College of Nursing Medicine and Health Sciences, University of Galway.

## CRediT authorship contribution statement

**Penglin Li:** Writing – review & editing, Writing – original draft, Visualisation, Methodology, Formal analysis, Conceptualisation. **Yating Zheng:** Writing – review & editing, Methodology, Formal analysis. **Jiwang Tang:** Methodology, Formal analysis. **Qin Xia:** Writing – review & editing, Methodology, Formal analysis. **José C. Casas-Martinez:** Writing – review & editing, Methodology, Formal analysis. **Ángel Ortiz-Alcántara**: Writing - review & editing, Methodology. **Raquel Requejo-Aguilar:** Writing - review & editing, Methodology. **C Alicia Padilla:** Writing - review & editing, Methodology. **Leo Quinlan**: Resources, Supervision, Formal Analysis. **Antonio Miranda-Vizuete**: Writing - review & editing, resources, Methodology: **Katarzyna Goljanek-Whysall:** Writing – review & editing, Supervision, Resources, Methodology. **Brian McDonagh:** Writing – review & editing, Writing – original draft, Supervision, Resources, Methodology, Formal analysis, Conceptualisation

