## Supplementary File for "Peroxiredoxin 6 limits mitochondrial peroxidation to prevent mitochondrial ER contact site assembly and inflammatory signalling following adaptive stress"

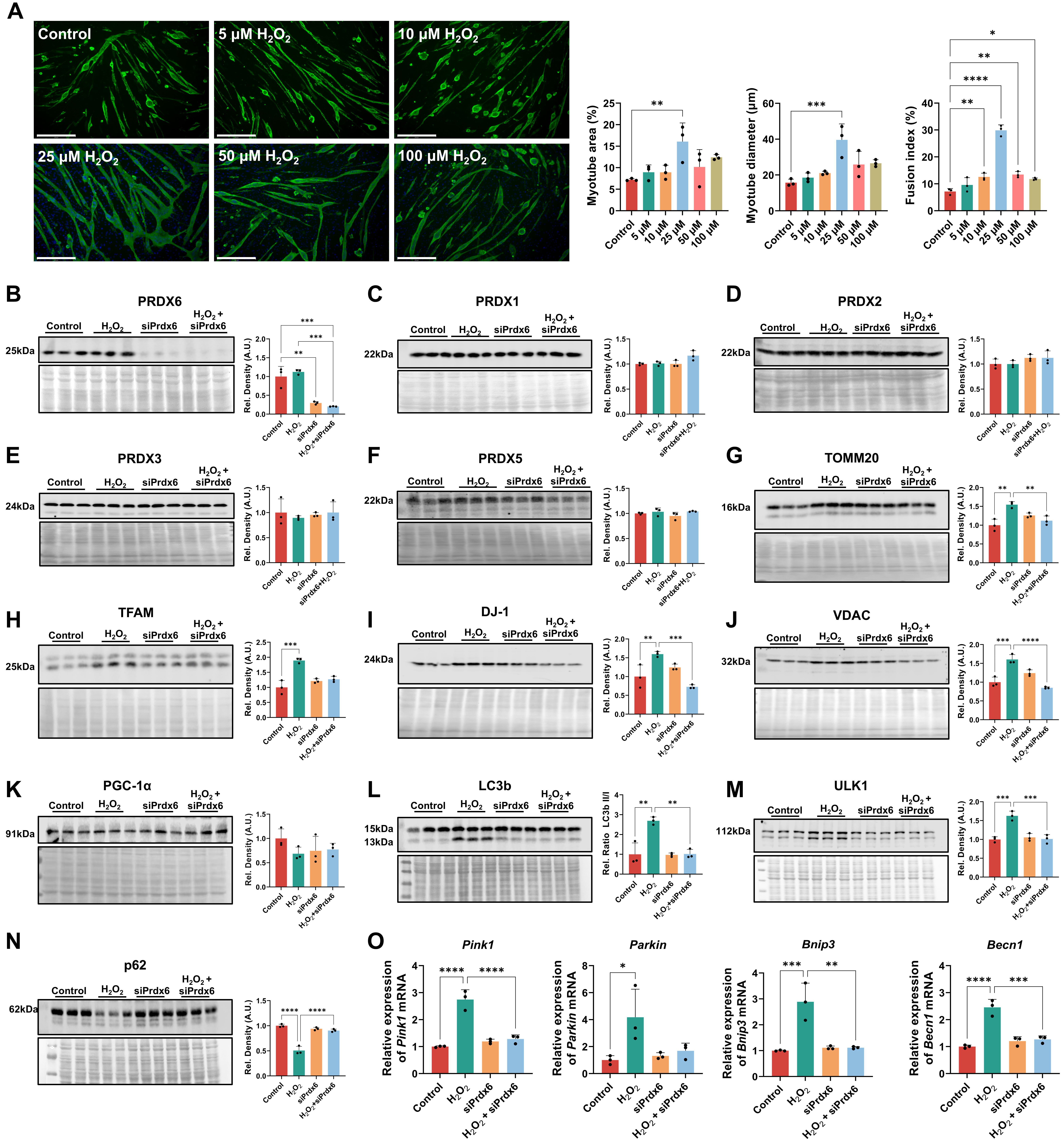

**Suppl. Fig.1 | The effects of H_2_O_2_ and Prdx6 knockdown on myogenesis, Peroxiredoxin levels, mitochondrial biogenesis, and mitophagy in myoblasts.** **(A)** Representative immunofluorescence images and quantification of myotubes after 7 days of differentiation following a 10 min treatment with H_2_O_2_ (0 to 100 µM). Scale bars, 275 µm. **(B-F)** Immunoblots and densitometry of Peroxiredoxins after control or siRNA knockdown of *Prdx6* and H_2_O_2_ treatment. B, PRDX6. C, PRDX1. D, PRDX2. E, PRDX3. F, PRDX5. **(G-K)** Immunoblots and densitometry of mitochondrial and biogenesis associated proteins. G, TOMM20. H, TFAM. I, DJ-1. J, VDAC1. K, PGC-1α. (**L-N),** Immunoblots and densitometry of autophagy markers. L**,** LC3B with the LC3 II to I ratio. M, ULK1. N, p62. **(O)** qPCR analysis of the mitophagy related genes *Pink1*, *Parkin*, *Bnip3* and *Becn1*. For all immunoblots, total protein was used as the loading control. Data presented as mean ± SEM. n = 3. For A, significance was determined by one-way ANOVA. For B-O, significance was determined by two-way ANOVA. * *p* < 0.05, ** *p* < 0.01, *** *p* < 0.001, **** *p* < 0.0001.

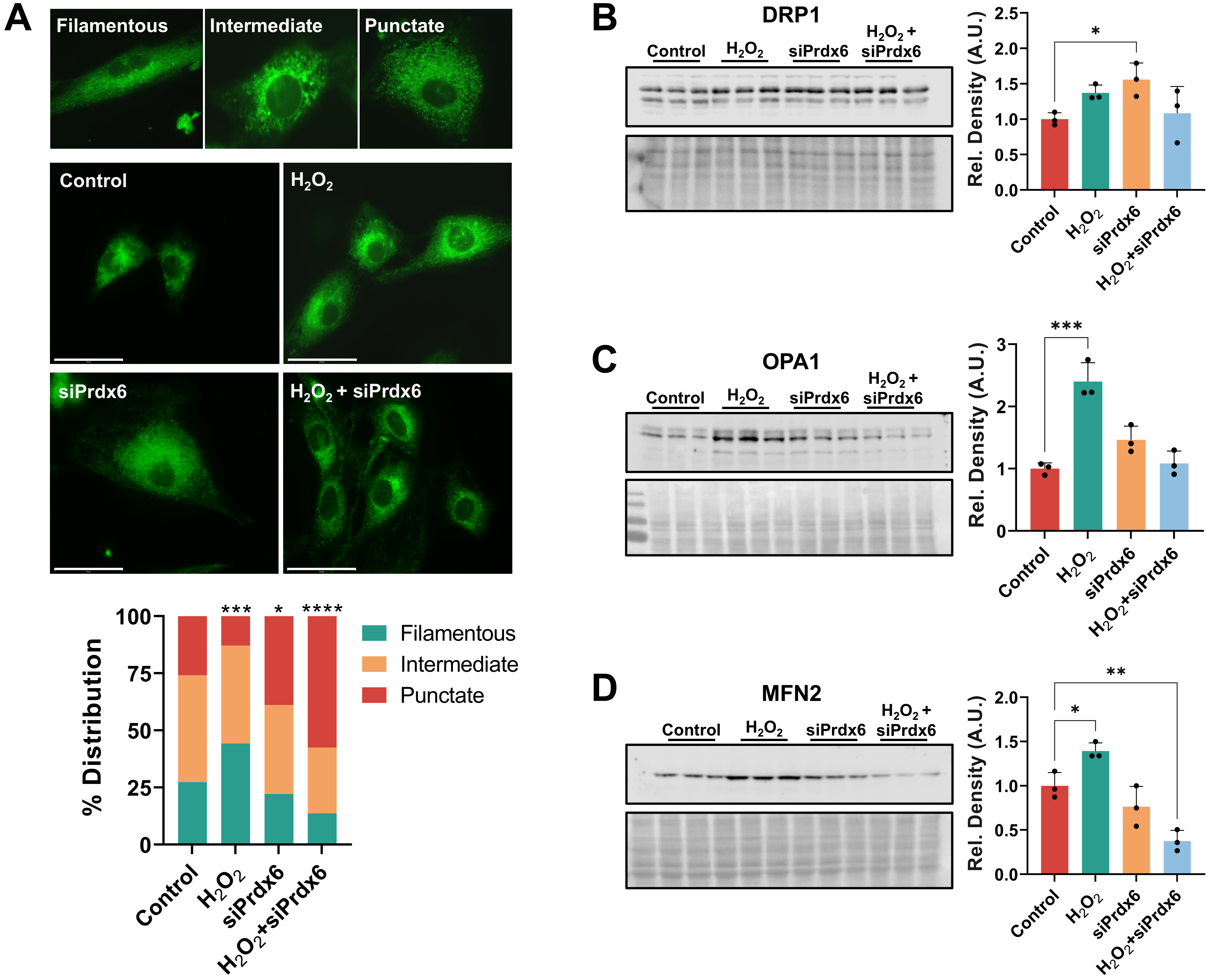

**Suppl. Fig.2 | Acute H_2_O_2_ promotes filamentous mitochondria in myoblasts that requires PRDX6. (A)** Mitochondrial network morphology after the 10 min treatment and recovery period. Top, representative images of the three scoring categories, filamentous, intermediate and punctate. Middle, representative images of myoblasts transfected with control or siRNA mediated knockdown on *Prdx6* and treated with 25 µM H_2_O_2_ or vehicle, with mitochondria labelled by MitoTracker Green. Bottom, stacked bar chart showing the percentage distribution of cells across the three morphology categories. Scale bars, 50 µm. **(B-D),** Immunoblots and densitometry of mitochondrial dynamics proteins DRP1, OPA1, and MFN2. Total protein was used as the loading control. For A, data represent the percentage distribution of 126-146 images per group; for B-D, n = 3 and data presented as mean ± SEM. Significance was determined by a chi-square test for the morphology distribution in A and by two-way ANOVA for B-D. **p* < 0.05, ** *p* < 0.01, *** *p* < 0.001, **** *p* < 0.0001.

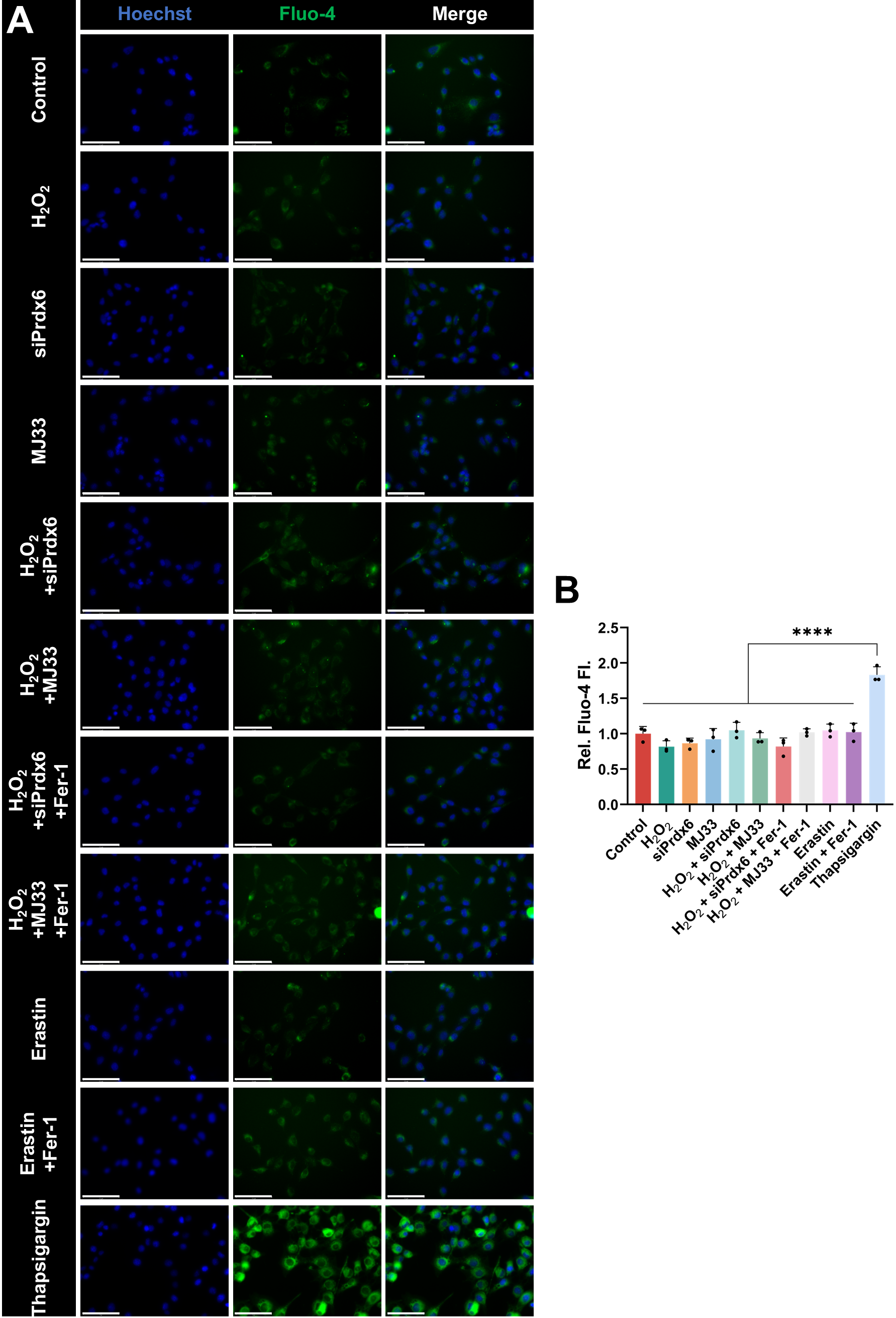

**Suppl. Fig.3 | Cytoplasmic calcium is unchanged under conditions that drive mitochondrial calcium influx.** Cytoplasmic Ca^2+^ was assessed by Fluo-4 staining in myoblasts. Thapsigargin was included as a positive control. (**A)** Representative images showing Hoechst, Fluo-4 and the merge. Scale bars, 50 µm. (**B)** Quantification of relative Fluo-4 fluorescence. Data presented as mean ± SEM. n = 3. Significance was determined by one-way ANOVA with Tukey’s multiple comparisons test. **** *p* < 0.0001.

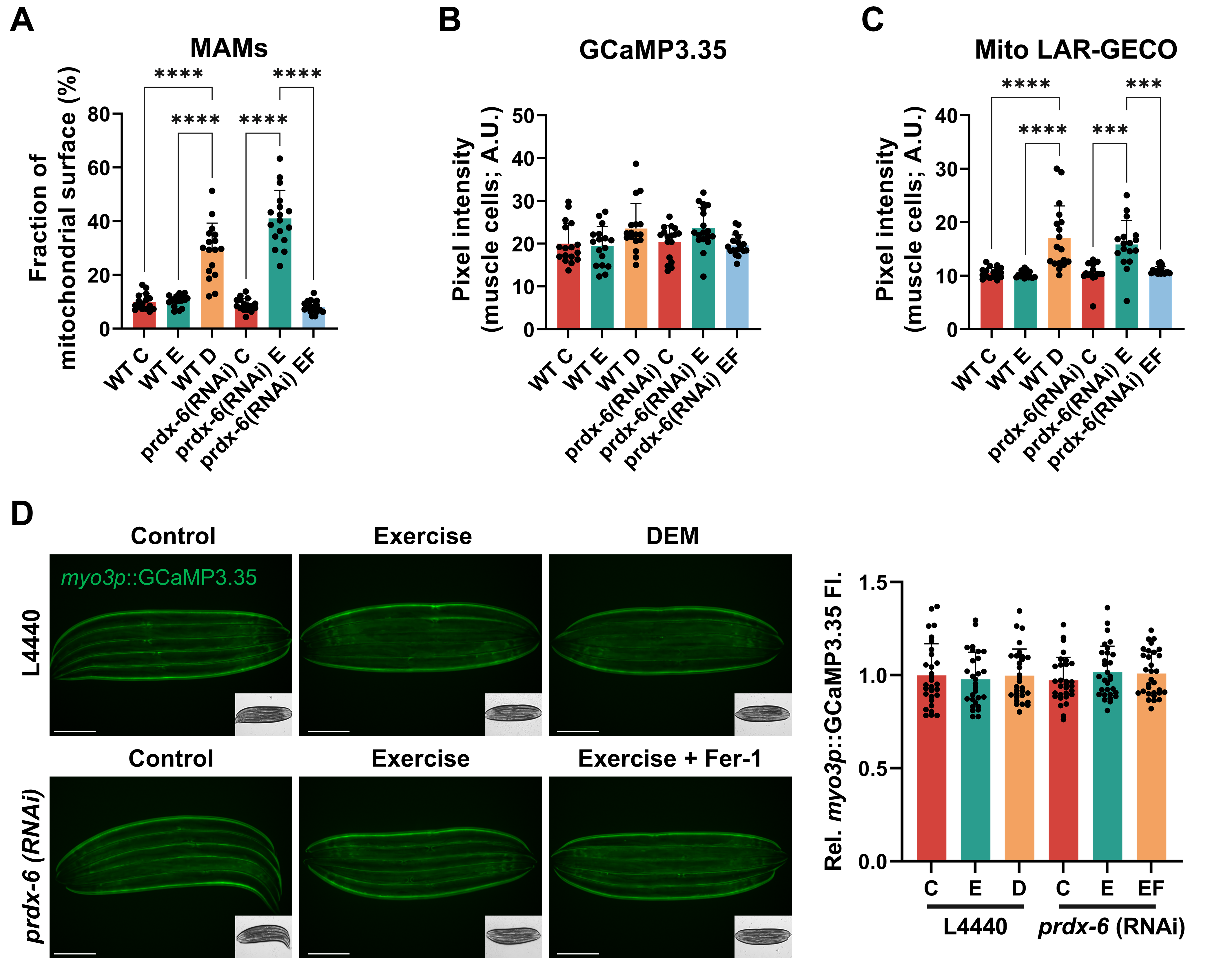

**Suppl. Fig.4 | Exercise promotes increased MAMs following exercise with RNAi mediated knockdown of *prdx-6* similar to wild type worms following ferroptosis induction with Diethyl maleate.** Animals were fed control (L4440) or *prdx-6* RNAi and subjected to the indicated treatments: control (C), exercise (E), diethyl maleate treatment (D), or exercise combined with ferrostatin-1 treatment (EF). (**A)** Quantification of the MAM images in Fig. 7I, expressed as the fraction of the mitochondrial surface in contact with the ER. **(B, C)** Quantification of the calcium images in Fig. 7J in muscle cells. B, Cytoplasmic calcium (GCaMP3.35). C, Mitochondrial calcium (mito LAR-GECO), each expressed as pixel intensity. **(D)** Total body wall muscle calcium assessed with the HBR4 reporter strain (*myo-3p*::GCaMP3.35). Representative images and quantification of relative *myo-3p*::GCaMP3.35 fluorescence. Scale bars, 275 µm. Data presented as mean ± SEM. n = 30. Significance was determined by one-way ANOVA *** *p* < 0.001, **** *p* < 0.0001.

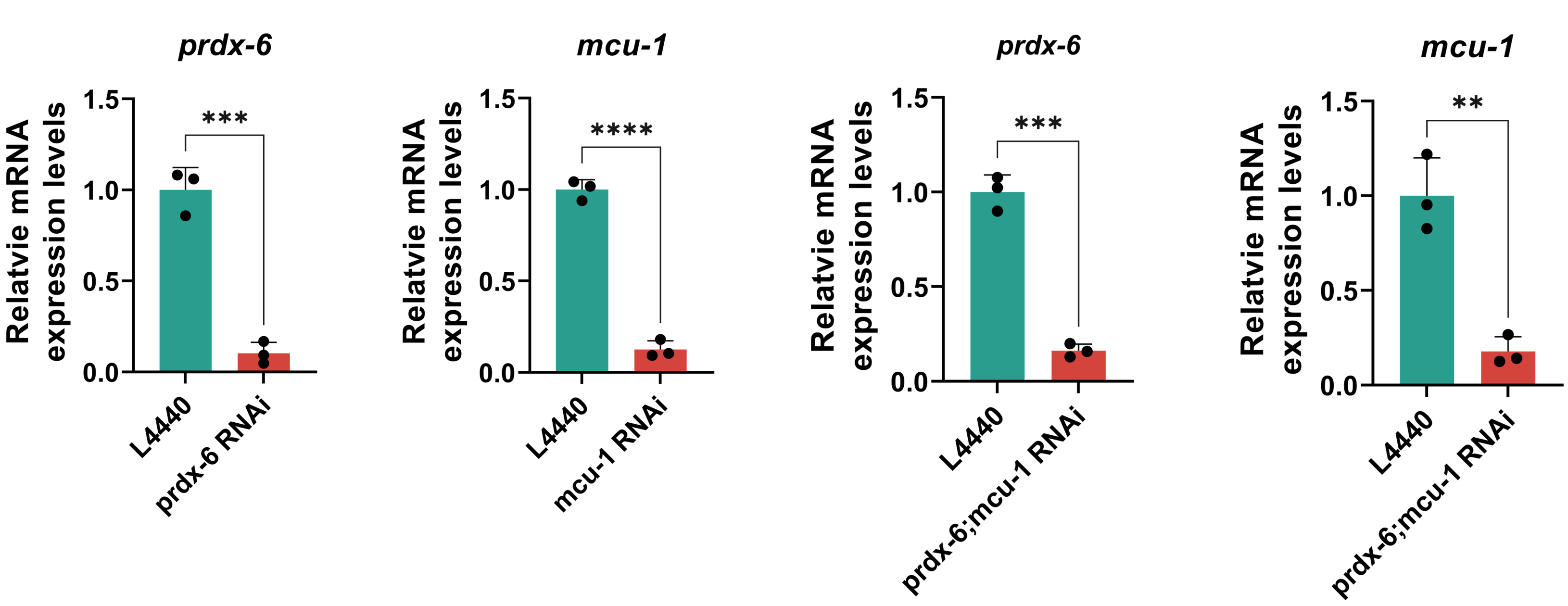

**Suppl. Fig.5 | Validation of RNA interference (RNAi) knockdown efficiency for target genes.** Relative mRNA expression levels of *prdx-6*, and *mcu-1* analysed by qPCR. Nematodes were fed E. coli HT115 expressing target-specific double-stranded RNA (dsRNA) or the L4440 empty vector control. Data represent the mean ± SEM (n = 3). ** *p* < 0.01, *** *p* < 0.001, **** *p* < 0.0001 compared to the L4440 control.

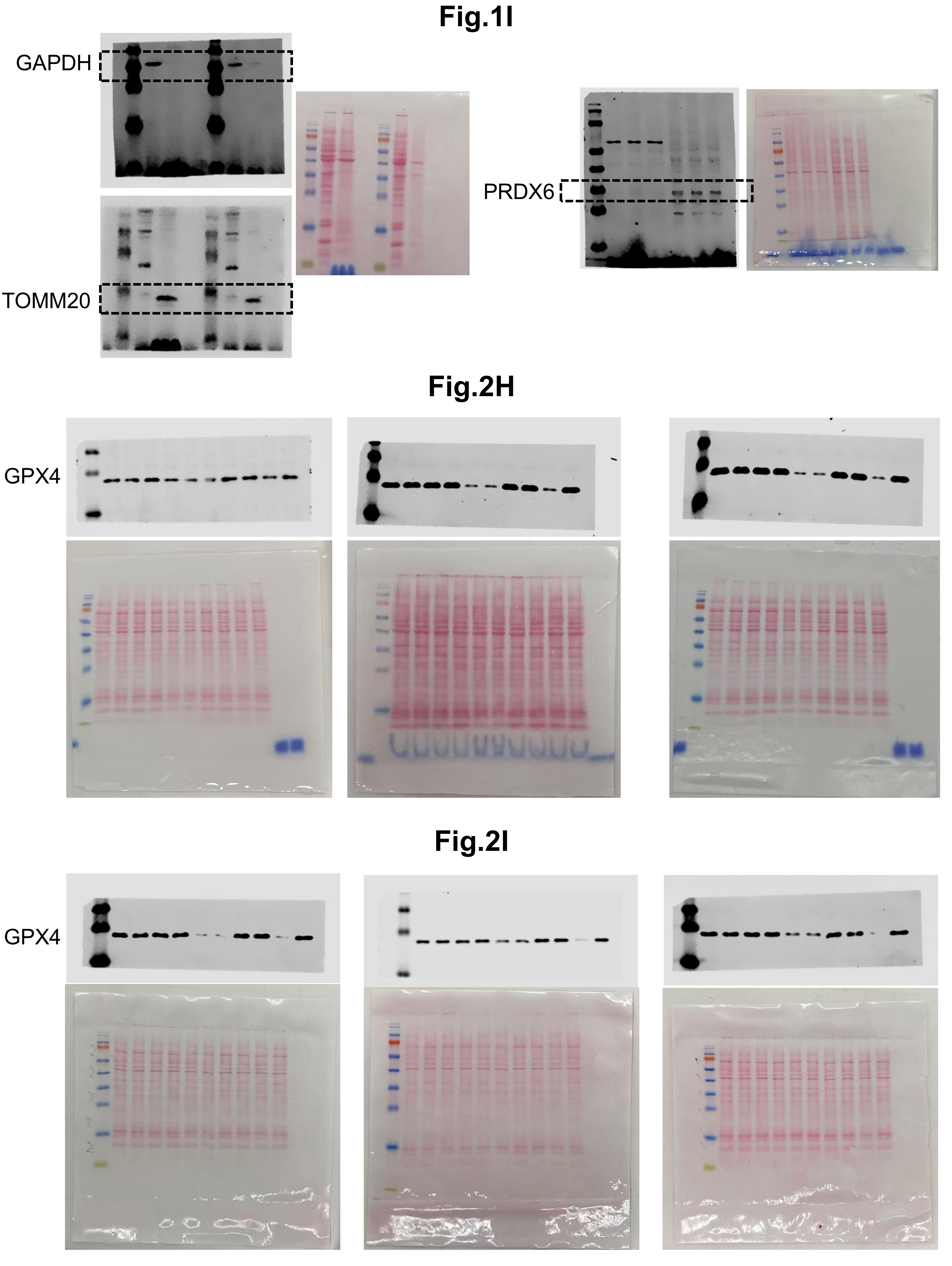

**Suppl. Fig.6 | Uncropped western blot images corresponding to Fig. 1I, 2H, and 2I.**

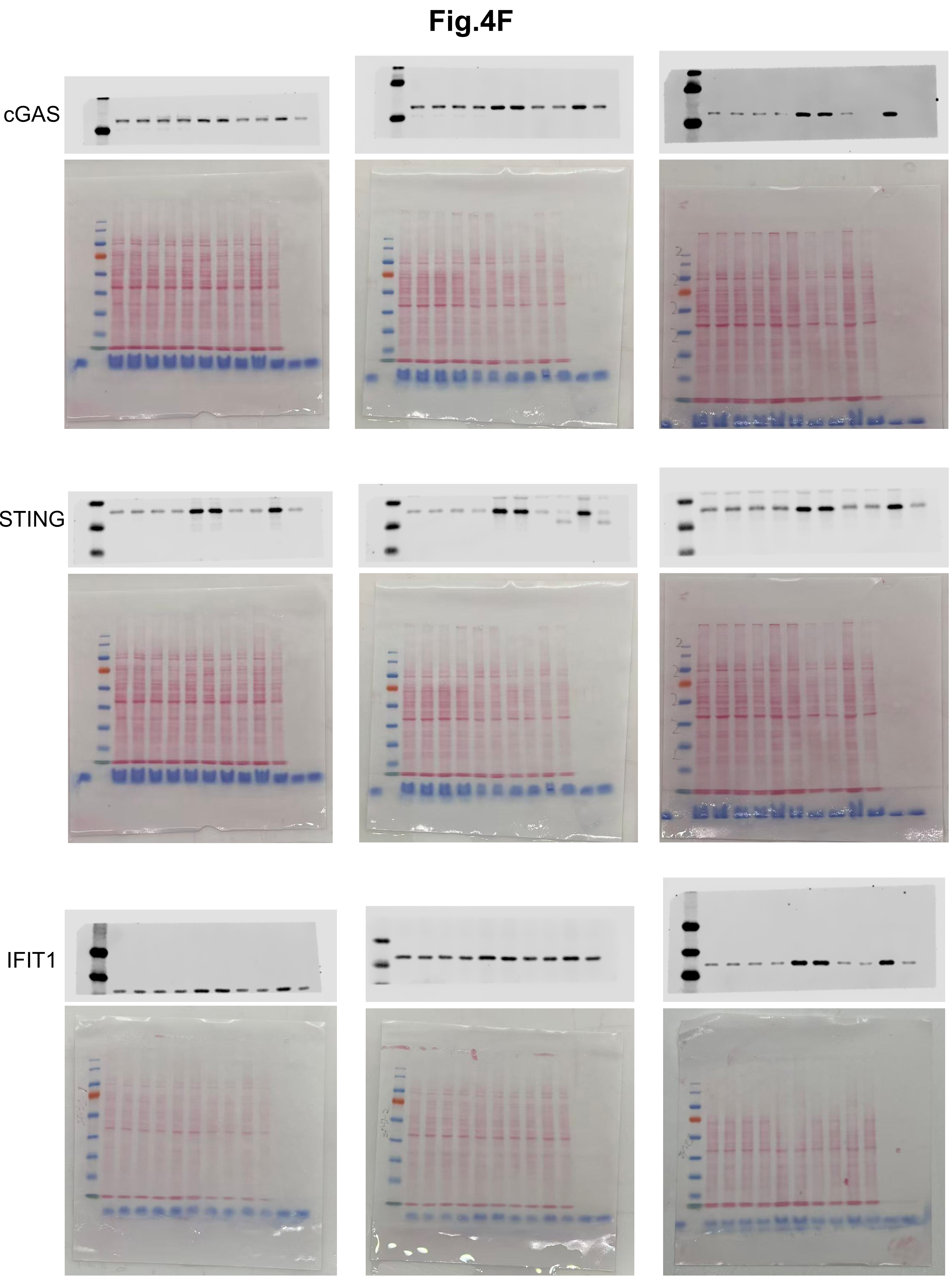

**Suppl. Fig.7 | Uncropped western blot images corresponding to Fig. 4F.**

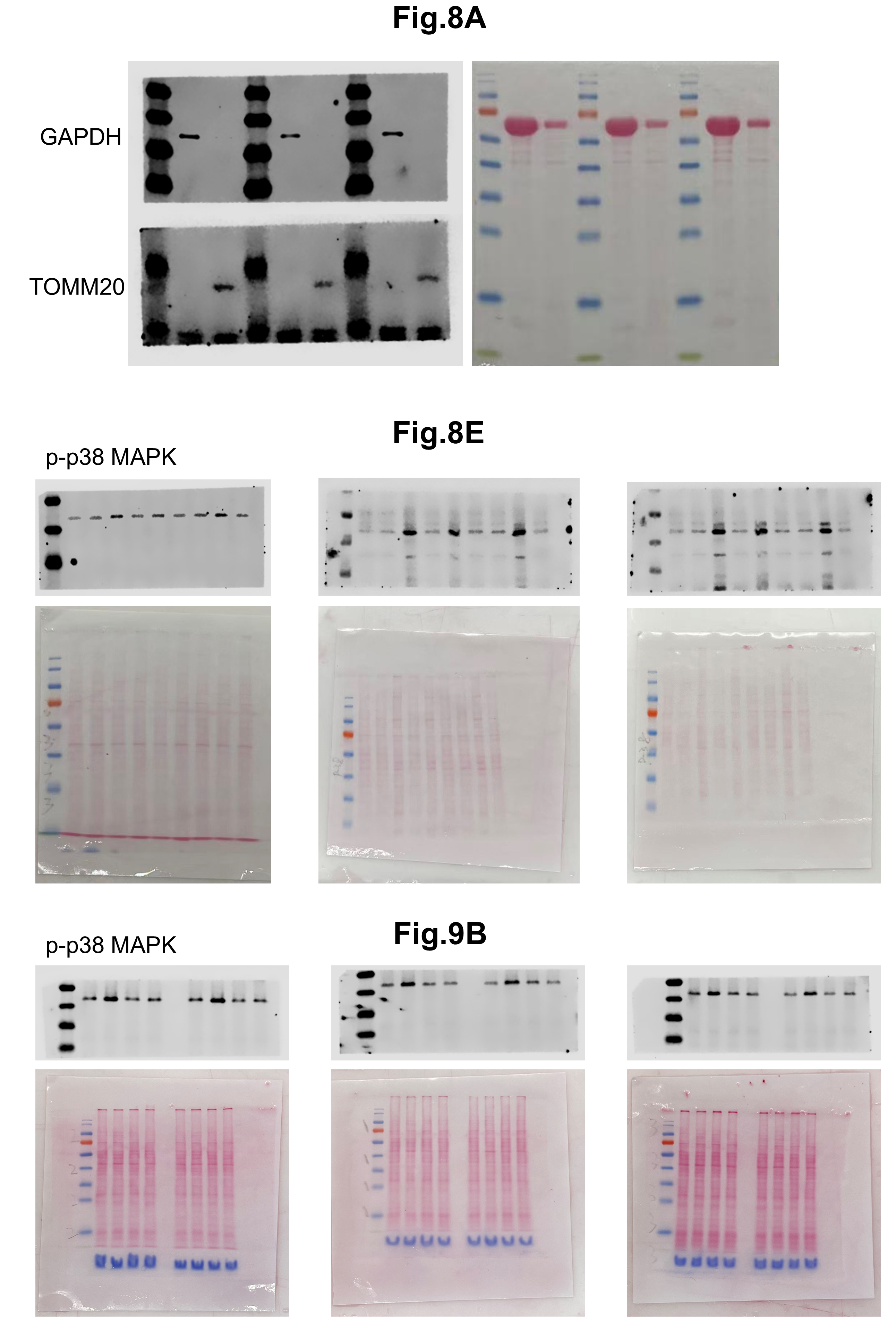
**Suppl. Fig.8 | Uncropped western blot images corresponding to Fig. 8A, 8E, and 9B.**

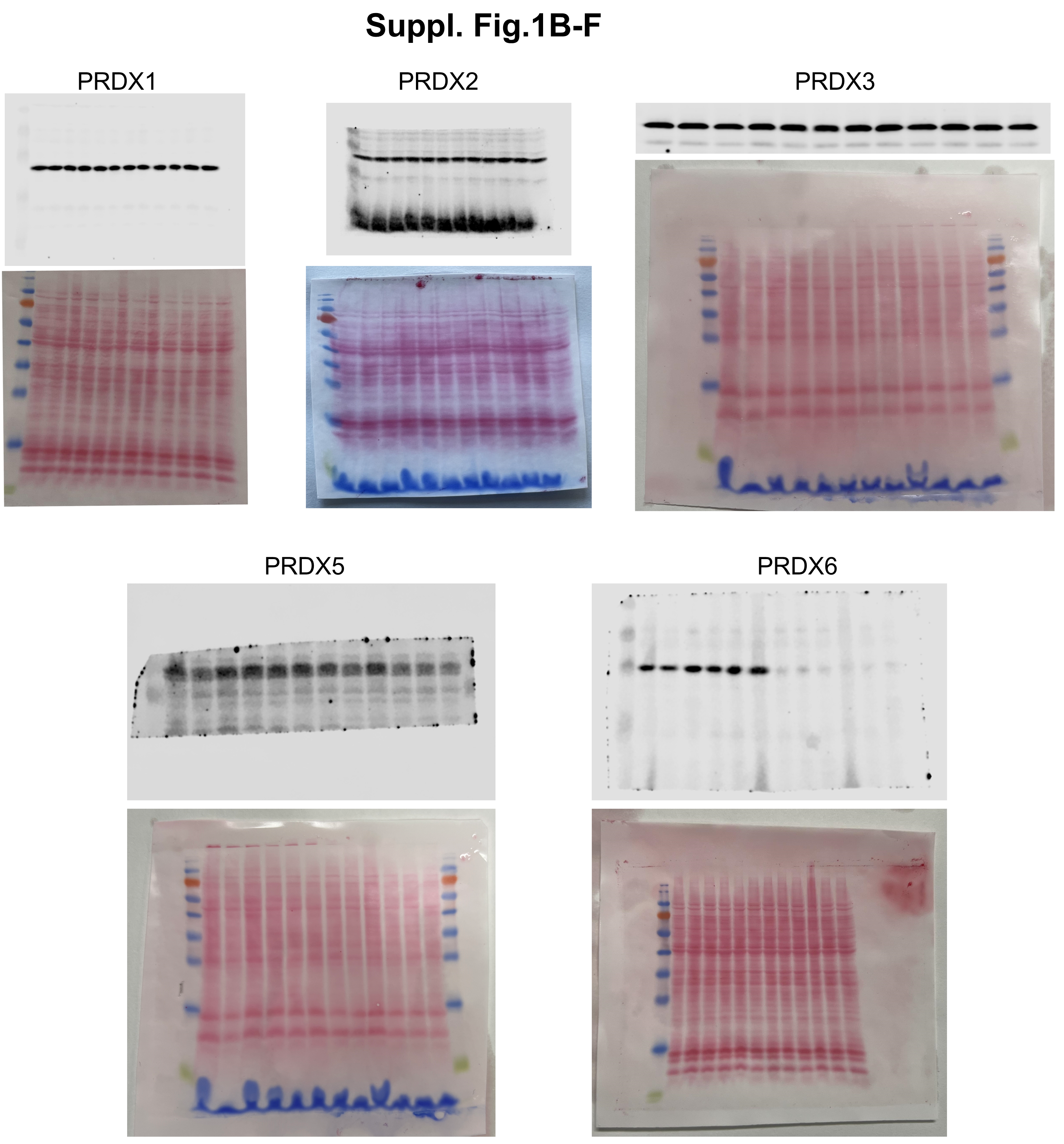

**Suppl. Fig.9 | Uncropped western blot images corresponding to Fig. 1B-F**

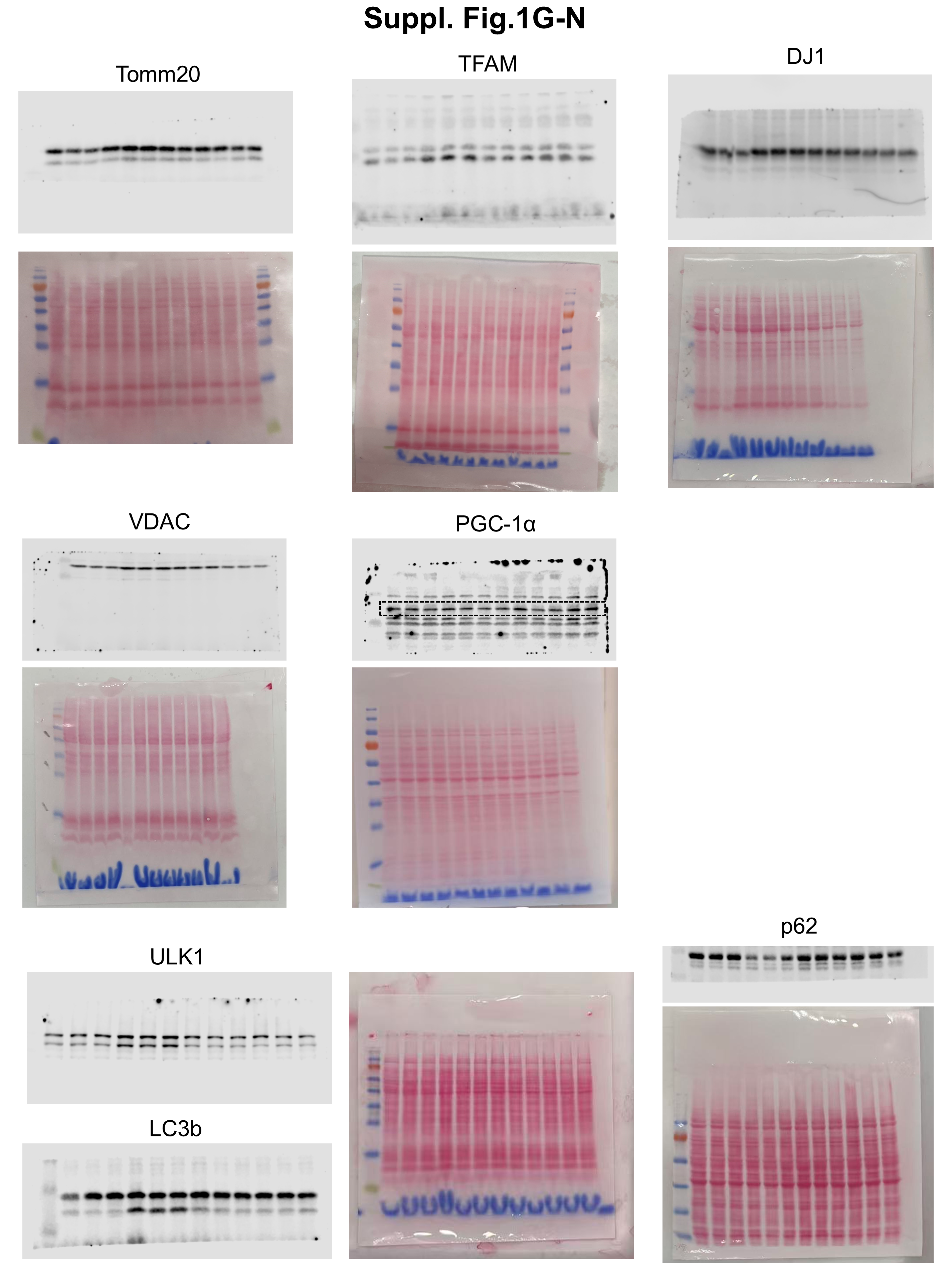

**Suppl. Fig.10 | Uncropped western blot images corresponding to Suppl. Fig. 1G-N.**

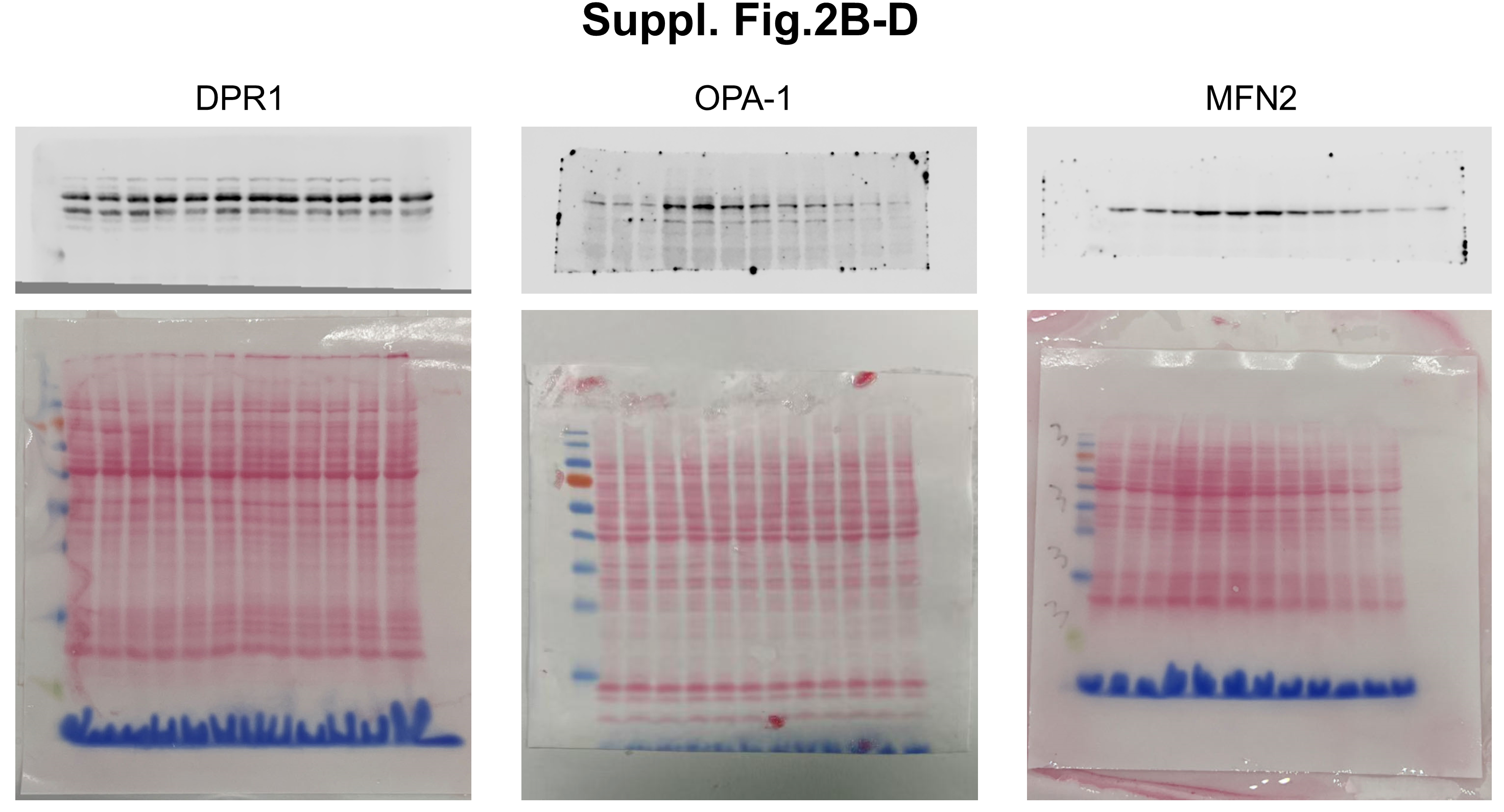

**Suppl. Fig.11 | Uncropped western blot images corresponding to Suppl. Fig.2B-D.**

**Supplementary Table 1 | Reagents and resources**

| **Cell line** | | |
| --- | --- | --- |
| C2C12 | ATCC | CRL-1772 |
| **Bacterial Strains** | | |
| E. coli: Strain OP50 | Caenorhabditis Genetics Center | N/A |
| *E. coli*: Strain HT115 (empty pL4440) | Miranda-Vizuete Lab | N/A |
| *E. coli*: Strain HT115 (RNAi clones) | Horizon Discovery (Cambridge, UK) | N/A |
| ***Caenorhabditis elegans* Strains** | | |
| Wild type | Caenorhabditis Genetics Center | N2 |
| *prdx-6*(*tm4225*) *IV* | NBRP *C. elegans* (Japan) | TM4225, Backcrossed 6X with wild type |
| *prdx-6*(*tm4284*) *IV* | NBRP *C. elegans* (Japan) | TM4284, Backcrossed 6X with wild type |
| *zcIs14[myo-3p::GFP(mit)]* | Caenorhabditis Genetics Center | SJ4103 |
| *unc-119(ed3); Ex[myo-3p::tomm20::Rosella; unc-119(+)]* | Tavernarakis Lab | IR2539 |
| ***unc-119(ed3); goeIs3[pmyo-3::GCaMP3.35::unc-54 3′UTR; unc-119(+)]*** | Caenorhabditis Genetics Center | HBR4 |
| *vkEx2674[pnhx-2CemOrange2::PISY-1; pmyo-2GFP]; zcIs17[pges-1mitGFP]* | Konstantinos Lab [46] | KPA427 |
| g*oeIs3[pmyo-3SL1::GCamP3.35::SL2; unc-119(+)]; aceIs1[pmyo-3mitoLAR-GECO; pmyo-2::RFP]* | Caenorhabditis Genetics Center | ATU2301 |
| *agIs44[irg-4p::GFP::unc-54–3*^′^*UTR; myo-2p::mCherry]* | Pukkila-Worley  Lab [50] | AU306 |
| *acIs101[irg-5p::GFP + rol-6(su1006)*]) | Caenorhabditis Genetics Center | AY101 |
| *agIs219[T24B8.5p::GFP::unc-54 3′UTR + ttx-3p::GFP::unc-54 3′UTR] III* | Caenorhabditis Genetics Center | AU78 |
| *prdx-6(tm4225) IV; zcIs14[myo-3p::GFP(mit)]* | This study | MCD27 |
| *prdx-6(tm4284) IV; zcIs14[myo-3p::GFP(mit)]* | This study | MCD28 |
| *prdx-6(tm4225) IV; unc-119(ed3); Ex[myo-3p::tomm-20::Rosella; unc-119(+)]* | This study | MCD30 |
| *prdx-6(tm4284) IV; unc-119(ed3); Ex[myo-3p::tomm-20::Rosella; unc-119(+)]* | This study | MCD33 |
| **Reagents and chemicals** | | |
| 2-Propanol | Merck | I9516 |
| 4′,6-diamidino-2-phenylindole (DAPI) | Merck | 268298 |
| Acetic acid | Sigma | A6283 |
| Acrylamide | Sigma | A3699 |
| Agar | Sigma | A1296 |
| Agarose Ultrapure | Thermo Fisher | 16500 |
| Ammonium acetate solution | Sigma | SLCM8842 |
| Ammonium Persulfate | Sigma | A3678 |
| Beta-mercaptoethanol | Sigma | M7522 |
| Bleach | Household | N/A |
| Bovine Serum Albumin BSA | Merck | A2153 |
| Bradford Reagent | Bio-Rad | 5000006 |
| Bromophenol blue | Sigma | 114391 |
| BSA | Sigma | A3059 |
| CaCl2 | Sigma | C1016 |
| Chloroform | Sigma | C0549 |
| Cholesterol | Sigma | C8667 |
| Digitonin | Sigma | D144 |
| DMEM | Sigma | D5796 |
| Erastin | Sigma | 329600 |
| Ethanol | Sigma | E7023 |
| Fast SYBR™ Green Master Mix | Applied | 4385617 |
| FBS | Sigma | F7524 |
| Ferrostatin 1 | TRC | TRC-F307700 |
| Fisher BioReagents™ EZ-Run™ Pre-stained Rec Protein Ladder | Fisher Bioreagents | BP36031 |
| Glucose | Sigma | G7021 |
| Glycogen | Merck | 10901393001 |
| Glycerol | Sigma | G6279 |
| Glycine | Sigma | G8898 |
| HEPES | Sigma | H3375 |
| H_2_O_2_ | Sigma | H1009 |
| Horse Serum | Thermo Fisher Scientific | 26050088 |
| Hydromount | Scientific laboratory supplies | D2176 |
| Isopropanol | Sigma | I9516 |
| KCl | Sigma | P3911 |
| K2HPO4 | Sigma | 3786 |
| KH2PO4 | Sigma | P9791 |
| Laminin | Merck | L2020 |
| Lipofectamine 2000 Reagent | Thermo Fisher Scientific | 11668019 |
| Methanol | Sigma | 34860 |
| MgCl2 | Sigma | M8266 |
| MgSO4 | Sigma | M7506 |
| MJ33 | Cayman chemical | 90001844 |
| N,N,N′,N′-Tetramethylethylenediamine | Merck | T9281 |
| Na2HPO4 | Sigma | S0876 |
| NaCl | Sigma | S9888 |
| NaOH | Sigma | S5881 |
| NEM | Sigma | E3876 |
| Nystatin | Sigma | N3503 |
| Paraformaldehyde | Thermo Fisher | 10131580 |
| Paraquat | Sigma | 856177 |
| PBS | Sigma | D8537 |
| Penicillin/Streptomycin | Thermo Fisher Scientific | 15140122 |
| Peptone | Sigma | 91249 |
| Phosphatase Inhibitor | Merck | P0044 |
| Ponceau S | Sigma | 141194 |
| Protease Inhibitor Cocktail | Sigma | P8340 |
| Random Hexamers (50 µM) | Thermo Fisher | N8080127 |
| RiboLock RNase Inhibitor (40U/µL) | Thermo Fisher | EO0381 |
| RNase A | Thermo Fisher | EN0531 |
| RNase free water | Sigma | W4502 |
| SDS | Sigma | L3771 |
| SuperScript™ II Reverse Transcriptase kit (includes 5X First-Strand Buffer and 0.1 M DTT) | Thermo Fisher | 18064014 |
| SYBR Green | Qiagen | 339347 |
| TEMED | Sigma | T9281 |
| Triton™ X-100 | Sigma | T8787 |
| Trizma® base | Sigma | T1503 |
| TRIzol Reagent | Life Technologies | 15596018 |
| TrypLE™ Express Enzyme | Thermo Fisher Scientific | 12604013 |
| Thapsigargin | Sigma | 586005 |
| Tween 20 | Merck | P9416 |
| **Antibodies** | | |
| Goat anti-Mouse IgG (H+L) Alexa Fluor™ Plus 488 | Thermo Fisher Scientific | A32727 |
| Goat anti-Mouse IgM (μ chain) Alexa Fluor™ 488 | Thermo Fisher Scientific | A-11035 |
| IRDye 800CW Goat anti-Mouse IgG | LI-COR Biosciences | 925-32210 |
| IRDye 800CW Goat anti-Rabbit IgG | LI-COR Biosciences | 926-32211 |
| Mouse anti-MF20 | Developmental Studies Hybridoma Bank | MF 20 |
| Mouse anti-DNA | Merck | CBL186 |
| Rabbit anti-MFN2 | Cell Signaling Technology | 9482 |
| Rabbit anti-OPA1 | Cell Signaling Technology | 80471 |
| Rabbit anti-DRP-1 | Cell Signaling Technology | 8570 |
| Rabbit anti-cGAS | Cell Signaling Technology | 31659 |
| Rrabbit anti-STING | Proteintech | 19851-1-AP |
| Rabbit anti-IFIT1 | Cell Signaling Technology | 14769 |
| Rrabbit anti-PGC-1alpha | Cell Signaling Technology | 2178 |
| Rabbit anti-TOM20 | Cell Signaling Technology | 42406 |
| Rabbit anti-Peroxiredoxin 1 | Abcam | ab109498 |
| Rabbit anti-Peroxiredoxin 2 | Cell Signaling Technology | 46855 |
| Mouse anti-Peroxiredoxin 3 | Abcam | ab16751 |
| Mouse anti-Peroxiredoxin 5 | Abcam | ab16944 |
| Rabbit anti-Peroxiredoxin 6 | Abcam | ab133348 |
| Rabbit anti-Phospho-p38 MAPK | Cell Signalling Technology | 4511 |
| Rabbit anti-TFAM | Abcam | ab272885 |
| Rabbit anti-ULK1 | Abcam | ab128859 |
| Rabbit anti-VDAC | Cell Signaling Technology | 4866 |
| Rabbit anti-LC3b | Abcam | ab192890 |
| Rabbit anti-SQSTM1/p62 | Cell Signaling Technology | 88588 |
| Rabbit anti-PARK7/DJ1 | Abcam | ab76241 |
| **Critical Commercial Assays** | | |
| C11-BODIPY581/591 | Merck | SML3717 |
| Fluo-4, AM | Thermo Fisher Scientific | F14201 |
| Malondialdehyde (MDA) Assay Kit | Solarbio life sciences | BC0025 |
| MitoTracker^TM^ Green FM | Thermo Fisher Scientific | M7514 |
| MitoSOX^TM^ Red mitochondrial superoxide indicator | Thermo Fisher Scientific | M36008 |
| MitoTracker^TM^ Deep Red FM | Thermo Fisher Scientific | M22426 |
| Rhod-2, AM | Thermo Fisher Scientific | R1245MP |
| TMRM | Thermo Fisher Scientific | I34361 |
| Glutathione (GSH) Assay Kit | Solarbio life sciences | BC1175 |
| **Plasmids** | | |
| pCLBW cox8 EGFP mCherry | Addgene | 78520 |
| ER-mit splitFAST | Riccardo Filadi Lab | N/A |
| **Primers for qRT-PCR** | | |
| Fw 5’-GGAGAATGGGAAGCCGAACA-3’ | Sigma | Mus_B2m |
| Rv 5’-TCTCGATCCCAGTAGACGGT-3’ | Sigma |  |
| Fw 5’-TCAGACAACTTTGGCCGACT-3’ | Sigma | Mus_Il-18 |
| Rv 5’-GGTGGATCCATTTCCACTTTGA-3’ | Sigma |  |
| Fw 5’-AAGCGGATGGACCTGGTGTTAGAACTG-3’ | Sigma | Mus_Ip3r1 |
| Rv 5’-AATTTGTGCTGTGTGCTTCGCGTAGAACT-3’ | Sigma |  |
| Fw 5’-TGAACGACGTGAAGACCTG-3’ | Sigma | Mus_Mcu |
| Rv 5’-CGAGCTCCCGCTCTTTGTTA-3’ | Sigma |  |
| Fw 5’-GTCTTTGCCGGAGAGCAGTA-3’ | Sigma | Mus_Camk1d |
| Rv 5’-GAGCTTCCTCTTCGGCAGTT-3’ | Sigma |  |
| Fw 5’-ACCCCACATGGGTTTGAGAC-3’ | Sigma | Mus_Ryr1 |
| Rv 5’-GACTCCTGACCAGTGTGCTC-3’ | Sigma |  |
| Fw 5’-GGCCCGAAACTACCTGGAGC-3’ | Sigma | Mus_Serca2a |
| Rv 5’-CAACGCACATGCACGCACCC-3’ | Sigma |  |
| Fw 5’-CTAGCTCATGTGTCAAGACCCTCTT-3’ | Sigma | Mus_Tert |
| Rv 5’-GCCAGCACGTTTCTCTCGTT-3’ | Sigma |  |
| Fw 5’-AATCTACCATCCTCCGTGAAACC-3’ | Sigma | Mus_D-loop |
| Rv 5’-TCAGTTTAGCTACCCCCAAGTTTAA-3’ | Sigma |  |
| Fw 5’-GCTTTCCACTTCATCTTACCATTTA-3’ | Sigma | Mus_Cytb |
| Rv 5’-TGTTGGGTTGTTTGATCCTG-3’ | Sigma |  |
| Fw 5’-CTAGAAACCCCGAAACCAAA-3’ | Sigma | Mus_Rnr2 |
| Rv 5’-CCAGCTATCACCAAGCTCGT-3’ | Sigma |  |
| Fw 5’-GAGGCTGTTGCTTGTGTGAC-3’ | Sigma | Mus_Nd1 |
| Rv 5’-TCTCGATCCCAGTAGACGGT-3’ | Sigma |  |
| Fw 5’-CTGATTACAAAAGAAGACATGACAGAC-3’ | Sigma | Mus_Ifi44 |
| Rv 5’-AGGCAAAACCAAAGACTCCA-3’ | Sigma |  |
| Fw 5’-CAAGGCAGGTTTCTGAGGAG-3’ | Sigma | Mus_Ifit1 |
| Rv 5’-GACCTGGTCACCATCAGCAT-3’ | Sigma |  |
| Fw 5’-CCAAGTGCTGCCGTCATTTTC-3’ | Sigma | Mus_Cxcl10 |
| Rv 5’-GGCTCGCAGGGATGATTTCAA-3’ | Sigma |  |
| Fw 5’-TCGAATGCGTTAGACTGCAC-3’ | Sigma | *cel_irg-4* |
| Rv 5’-TAGCACTGGTTCCTTGGCTC-3’ | Sigma |  |
| Fw 5’-CATGGACCTGATTACACCGCT-3’ | Sigma | *cel_irg-5* |
| Rv 5’-TCGTACTTCTTCACCGCAGAT-3’ | Sigma |  |
| Fw 5’-AGACCATCATGCCCTTCACT-3’ | Sigma | *cel_t24B8.5* |
| Rv 5’-GTAACGCAGACACCACAGGT-3’ | Sigma |  |
| Fw 5’-ATGCTACCGAATGCTTTTCC-3’ | Sigma | *cel_k08D8.5* |
| Rv 5’-TCCTTGGGTGTAGTTTCCAA-3’ | Sigma |  |
| Fw 5’-GGATACTAATGGACCAACTACATT-3’ | Sigma | *cel_k08D8.4* |
| Rv 5’-TCCATCTTGTCTCCCAAGATAC-3’ | Sigma |  |
| Fw 5’-CGAAGCCAACAACGGAAAGTA-3’ | Sigma | *cel_tbb-2* |
| Rv 5’-TCCGAACACAAAGTTGTCAGG-3’ | Sigma |  |
| Fw 5’-GTTATTGCAGTGCCAACAGGT-3’ | Sigma | *cel_cox-1* |
| Rv 5’-ACAACACCTGTCAACCCACC-3’ | Sigma |  |
| Fw 5’-GGGATGTTGGTGACATTGCC-3’ | Sigma | *cel_cytb-1* |
| Rv 5’-TGCTATTAACCTATCGGGCGT-3’ | Sigma |  |
| Fw 5’-TAACCGGGCGCCATTTGATT-3’ | Sigma | *cel_nd-1* |
| Rv 5’-GCTACTCTGGCAAACTCCACA-3’ | Sigma |  |
| Fw 5’-ATCCTATTCACAACGCCCGA-3’ | Sigma | *cel_aat-9* |
| Rv 5’-ACGACAGATTCATCGTCGGATT-3’ | Sigma |  |
| Fw 5’-CGAAGCGGCCGTCAATAAAC-3’ | Sigma | *cel_ftn-1* |
| Rv 5’-GGGCCGGCTCTCTTGATATT-3’ | Sigma |  |
| Fw 5’-GTGATGACGTGTCACTTTCGG-3’ | Sigma | *cel_gpx-1* |
| Rv 5’-AAGTCCACACTGTGAAGCGA-3’ | Sigma |  |
| Fw 5’-GTCGGAGAGAGTCAAGGCTG-3’ | Sigma | *cel_acs-17* |
| Rv 5’-ACGTCGGACACATTCTTCCC-3’ | Sigma |  |
| Fw 5’-CAGAGTTCGTTCCCGAGCAT-3’ | Sigma | *cel_itr-1* |
| Rv 5’-GTGTGCTTCCACCACCACTA-3’ | Sigma |  |
| Fw 5’-TGAGCGACGTTTTTAGTCTGTT-3’ | Sigma | *cel_unc-68* |
| Rv 5’-ATCACCCAAAGTGGTTCGCT-3’ | Sigma |  |
| Fw 5’-CGACTTCGGAGCTTCCCTTT-3’ | Sigma | *cel_vdac-1* |
| Rv 5’-ACAGTAAGGTCACGGCTTGG-3’ | Sigma |  |
| Fw 5’-TTCAACTGGAGAACGCCGAA-3’ | Sigma | *cel_mcu-1* |
| Rv 5’-ACTCGATCCGTGTGAGCTTC-3’ | Sigma |  |
| Fw 5’-AGCCATTCTGGCCGCTCTCG-3’ | Sigma | *cel_cdc-42* |
| Rv 5’-GCAACCGCTTCTCGTTTGGC-3’ | Sigma |  |
| **Software** | | |
| Prism 10.5 | GraphPad Software | RRID:SCR_002798 |
| ImageJ | NIH | RRID:SCR_003070 |
| Image Studio Lite | Image Studio Lite | RRID:SCR_013715 |
| CeleST | Restif et al., 2014 [65] | N/A |
| OASIS 2 | Online Application for Survival Analysis 2 | N/A |

**Supplementary Table 2 | Data from survival assays**

| **Figure** | **Group**  **(Strain and condition)** | **Deaths/ Total** | **Mean survival** | **Log-rank test, *p*-value (Compared to control)** |
| --- | --- | --- | --- | --- |
| 5A | N2C | 101/105 | 14.64 ± 0.38 | 0.0002$ |
|  | N2E | 102/105 | 16.63 ± 0.39 |  |
|  | *tm4225* C | 99/105 | 12.83 ± 0.35 | 0.0080$ |
|  | *tm4225* E | 100/105 | 11.59 ± 0.31 |  |
|  | *tm4284* C | 103/105 | 13.03 ± 0.38 | 0.0051$ |
|  | *tm4284* E | 100/105 | 11.70 ± 11.7 |  |
| 5B | N2 C | 50/50 | 25.68 ± 2.29 | 0.0206$ |
|  | N2 E | 50/50 | 31.80 ± 2.56 |  |
|  | *tm4225* C | 50/50 | 27.24 ± 1.65 | 0.0151$ |
|  | *tm4225* E | 50/50 | 24.06 ± 1.20 |  |
|  | *tm4284* C | 50/50 | 31.24 ± 1.85 | < 0.0001$ |
|  | *tm4284* E | 50/50 | 20.80 ± 1.56 |  |
| 5C | N2 C | 50/50 | 11.5 ± 0.95 | < 0.0001$ |
|  | N2 E | 50/50 | 16.83 ± 1.22 |  |
|  | *tm4225* C | 50/50 | 11.84 ± 0.58 | 0.0038$ |
|  | *tm4225* E | 50/50 | 9.68 ± 0.47 |  |
|  | *tm4284* C | 50/50 | 12.24 ± 0.53 | 0.0053$ |
|  | *tm4284* E | 50/50 | 9.96 ± 0.49 |  |
| 9A | *tm4225*;L4440 C | 105/105 | 14.24 ± 0.34 | 0.0063* |
|  | *tm4225*;L4440 E | 104/105 | 13.12 ± 0.29 |  |
|  | *tm4225*;mcu-1(RNAi) C | 104/105 | 14.21 ± 0.33 |  |
|  | *tm4225*;mcu-1(RNAi) E | 105/105 | 14.59 ± 0.34 | 0.0002# |
|  | *tm4225*;L4440; Fer-1 C | 105/105 | 14.42 ± 0.33 |  |
|  | *tm4225*;L4440; Fer-1 E | 102/105 | 15.63 ± 0.36 | < 0.0001# |
|  | *tm4225*;mcu-1(RNAi); Fer-1 C | 103/105 | 14.55 ± 0.35 |  |
|  | *tm4225*;mcu-1(RNAi); Fer-1 E | 103/105 | 15.38 ± 0.35 | < 0.0001# |
|  | *tm4284*;L4440 C | 105/105 | 14.10 ± 0.33 | 0.0272* |
|  | *tm4284*;L4440 E | 104/105 | 13.01 ± 0.27 |  |
|  | *tm4284*;mcu-1(RNAi) C | 104/105 | 14.25 ± 0.33 |  |
|  | *tm4284*;mcu-1(RNAi) E | 104/105 | 14.63 ± 0.35 | < 0.0001# |
|  | *tm4284*;L4440; Fer-1 C | 105/105 | 14.49 ± 0.34 |  |
|  | tm4284;L4440; Fer-1 E | 105/105 | 15.33 ± 0.36 | < 0.0001# |
|  | *tm4284*;mcu-1(RNAi); Fer-1 C | 104/105 | 14.45 ± 0.33 |  |
|  | *tm4284*;mcu-1(RNAi); Fer-1 E | 104/105 | 15.46 ± 0.35 | < 0.0001# |
| $ (Compared to the control in each strain)  * (Compared to *tm4225*;L4440 C / *tm4284*;L4440 C)  # (Compared to *tm4225*;L4440 E / *tm4284*;L4440 E) | | | | |
